# Cortical Mechanisms for Reaches Versus Saccades: Progression of Effector-Specificity Through Target Memory to Movement Planning and Execution

**DOI:** 10.1101/415562

**Authors:** David C. Cappadocia, Simona Monaco, Ying Chen, J. Douglas Crawford

## Abstract

Effector-specific cortical mechanisms can be difficult to establish using fMRI, in part because low time resolution might temporally conflate different signals related to target representation, motor planning, and motor execution. Here, we used an event-related fMRI protocol and a cue-separation paradigm to temporally separate these three major sensorimotor stages for saccades vs. reaches. In each trial, subjects (N=12) 1) briefly viewed a target 4-7° left or right of midline fixation on a touchscreen, followed by an 8 second delay (effector-independent *target memory* phase), 2) were instructed by an auditory cue to perform a reach or a saccade, followed by a second delay of 8 seconds (effector-specific *planning* phase), and finally 3) were prompted to move by reaching-to-touch or performing a saccade towards the remembered target (effector-specific *execution* phase). Our analysis of saccade and reach activation (vs. a non-spatial control task) revealed modest effector-agnostic target memory activity (left AG, bilateral mIPS) followed by independent effector parietofrontal sites and time courses during the motor components of the task, specifically: more medial (pIPS, mIPS, M1, and PMd) activity during both *reach planning* and *execution*, and more lateral (mIPS, AG, and FEF) activity only during *saccade execution*. These motor activations were bilateral, with a left (contralateral) preference for reach. A conjunction analysis revealed that left mIPS and right AG, PCu, SPOC, FEF/PMv and LOTC showed activation for both saccades and reaches. Overall, effector-preference contrasts (reach vs. saccade) revealed significantly more parietofrontal activation for reaches than saccades during both planning and execution, with the exception of FEF. Cross-correlation of reach, saccade, and reach-saccade activation through time revealed correlated activation both within and across effectors in each hemisphere, but with a tendency toward higher correlations in the right hemisphere, especially *between* the eye and hand. These results demonstrate substantially independent but temporally correlated cortical networks for human eye, hand, and eye-hand control, that follow explicit spatiotemporal rules for effector-specific timing, medial-lateral distribution, and hemispheric lateralization.

## Introduction

Human beings rely on the input of sensory information to guide actions and effectively interact with their environment. Two of the most frequent goal-directed actions that use visual information are visually guided saccades and reaches (e.g., a car driver moving their eyes from the road to their rear-view mirror or their right hand from the steering wheel to the radio). In monkeys, very specific signals have been localized for saccades versus reaches in frontal and parietal cortex (Buneo and Andersen, 2002; Synder et al., 2000). In contrast, reach and saccade signals have been very difficult to dissociate in human functional magnetic resonance imaging (fMRI) studies of parietal cortex (Vesia and Crawford 2012). Indeed, several studies have suggested highly distributed reach regions (Filimon et al., 2007; 2009) with large amounts of overlap with saccade activity during planning and execution (Medendorp et al., 2003; Beurze et al., 2007, 2009). These distributed responses have resulted in considerable controversy regarding the localization and degree and of effector specificity in human cortex.

To briefly summarize fMRI evidence for saccade-reach effector specificity in posterior parietal cortex (PPC), a cluster of activation in the midposterior intraparietal sulcus (mIPS) has been linked to saccade planning (Astafiev et al., 2003; Merriam et al., 2003; Schluppeck et al., 2005; Tosoni et al., 2008), eye movements and attention (Corbetta et al., 1998; Astafiev et al., 2003), eye movements and visual working memory (Curtis et al, 2004; Curtis and Connelly, 2008; Srimal and Curtis, 2008) and all three of these processes (Jerde et al., 2012). In reach, human mIPS has been linked to reach planning (DeSouza et al., 2000; Medendorp et al., 2003, 2005; Prado et al., 2005; Beurze et al., 2007, 2009, 2010; Fernandez-Ruiz, 2007; Tosoni et al., 2008; Filimon et al., 2009; Chen et al., 2014; Cappadocia et al., 2017), as have other parietal regions including the superior parietal occipital cortex (SPOC) (Astafiev et al., 2003; Connolly et al., 2003; Prado et al., 2005; Fernandez-Ruiz et al., 2007; Tosoni et al., 2008; Beurze et al., 2009; Gallivan et al., 2009, 2011; Bernier and Grafton, 2010; Cavina-Pratesi et al., 2010; Monaco et al., 2011; Chen et al., 2014; Cappadocia et al., 2017) and the angular gyrus (AG) (Fernandez-Ruiz et al., 2007; Chen et al., 2014; Cappadocia et al., 2017). In frontal cortex, several studies have examined saccade related activation in the human frontal eye fields (FEF) (Astafiev et al., 2003; Beurze et al., 2009; Amiez and Petrides, 2009; Gallivan et al, 2011; Herwig et al, 2014), and reach related activation in the human dorsal premotor cortex (PMd) (Connolly et al, 2007; Curtis et al, 2008; Gallivan et al, 2011, 2015; Bestman et al., 2012).

Despite this progress, several questions remain unresolved. Many of the fMRI studies cited above reported considerable overlap between saccade and reach activity, either due to actual sharing of control, sharing of common inputs required for both saccades and reach, or simply due to the spatial resolution limits of fMRI (Connolly et al, 2007; Curtis et al, 2008; Beurze et al., 2009; Gallivan et al, 2011). Specific questions include the extent to which reach and saccade signals are segregated medially versus laterally relative to the intraparietal sulcus (Culham et al., 2006; Gallivan et al., 2011). Also, there is emerging evidence that reach activation is lateralized to the hemisphere contralateral to the hand as early as parietal and even occipital cortex (Fernandez-Ruiz et al., 2007; Cappadocia et al., 2017). Since the two eyes are normally yoked one would not expect to see the same type of mass contralateral lateralization during saccades, but the degree of general lateralization, if any, for saccades in humans remains unclear (Chen et al. 2016).

One factor that has not been fully considered in this debate is how the low *temporal* resolution of fMRI might interact with multiple neural signals to conflate the spatial resolution of effector-independent, effector-dependent (reach or saccade related activation), and effector-specific (activation specific to reach or saccade and not the other) signals. Specifically, studies that do not separate target and memory responses from motor planning could yield ‘planning’ activation that is actually a combination of these three types of activation. Conversely, studies that do not separate planning from execution might confuse spatial specificity with effector independence if the temporal distribution of activation through these phases is different for saccade and reach. In short, these paradigms might tend to conflate different signals and thus underestimate the spatial specificity of motor effector signals (Cappadocia et al., 2017).

The current study exploited a recently developed cue-separation paradigm (Cappadocia et al., 2017) to examine the temporal progression of neural correlates of both independent and preferential planning and execution for reach and saccades. This study uses an event-related fMRI paradigm that explicitly separates visually-guided reaches and saccades into three phases in time (effector agnostic visual target representation, independent effector movement planning, and independent effector movement execution), by introducing an effector instruction between visual target memory and planning phases, and a ‘go signal’ between planning and execution times. We then analyzed the concomitant BOLD activation in cortex to investigate how independent and preferential effector-specific planning and execution signals are temporally and spatially distributed through the cortical networks for action in the human, and the degree to which these signals are correlated through time within and across the reach and saccade networks. We find that, when visuospatial activation is disentangled from motor activation, the cortical networks for human reach and saccade control show substantially different distributions in both the spatial and temporal domains.

## Methods

### Participants

Twelve right-handed subjects (3 males, 9 females aged 20-36) were recruited from the York University community. We chose this number of subjects based on precedents set in similar studies of visuomotor control in healthy subjects (Cavina-Pratesi et al., 2007, Gallivan et al., 2011). The resulting dataset was sufficient to yield statistically significant results that survived corrections for multiple comparisons (see Results). All subjects had normal or corrected-to-normal vision and none of the subjects had any known neurological deficits. The York University Human Participants Review Sub-committee approved all techniques used in this study and all participants gave their informed consent prior to the experiment.

### Experimental stimuli and apparatus

The experimental stimuli and apparatus were the same as the setup used in Chen et al. (2014) and Cappadocia et al. (2017). Visual stimuli consisted of optic fibers embedded into a custom-built board with adjustable tilt. The board was mounted atop a platform whose height was also adjustable (Figure 1A). The platform was attached to the MRI scanner bed and placed over the abdomen of the subject. The height of the platform and tilt of the board were adjusted for each participant to ensure comfortable reaching movements. A translucent touchscreen (Keytec, 170 mm X 126 mm) was affixed on the board to record reach endpoints. An eye-tracking system (iView X) was used in conjunction with the MRI-compatible Avotec Silent Vision system (RE-5701) to record movements of the right eye during the experiment.

**Figure 1.**
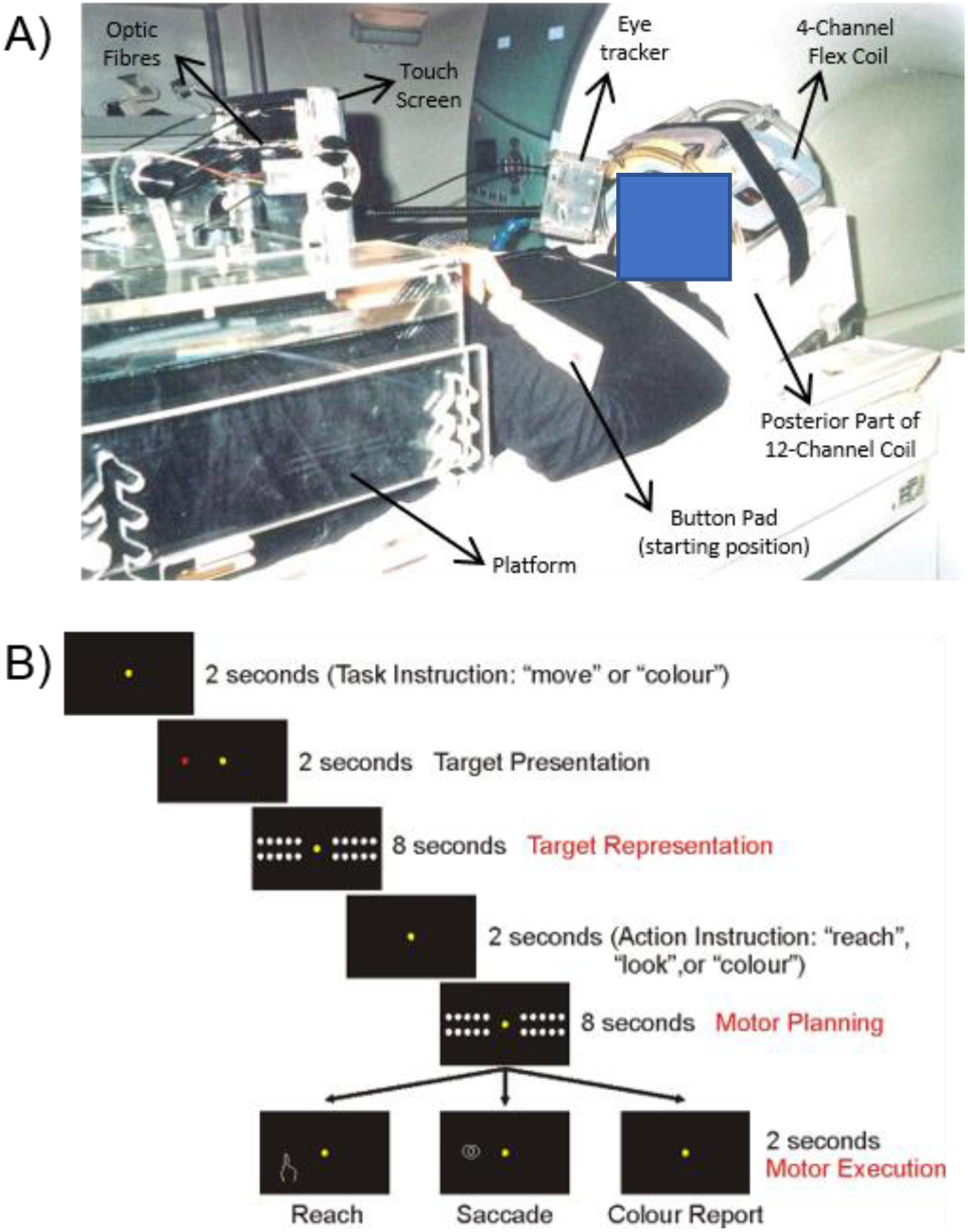
Experimental setup and paradigm. **A**) Photograph of The experimental setup. ***B)*** Illustration of the experimental paradigm. The display of visual targets is the same for all three tasks (Reach, Saccade, and Color Report). The key difference between the two action tasks is the auditory effector command, informing the effector to be used. As the target presentation and effector instruction are separated by an 8 second delay, this allows the task to disentangle target representation from movement planning and execution. In the Color Report task, target color (red or green) rather than location is remembered and reported.

The head of the participant was slightly tilted (∼20°) to allow direct viewing of the stimuli presented on the board (Figure 1A). The board was approximately perpendicular to gaze and ∼60 cm from the eyes. The upper arm was strapped to the scanner bed to limit motion artifacts during reach trials. Reaches were thus performed by movements of the right forearm and hand. A button pad was placed on the left side of the participants’ abdomen and served as both the starting point for each trial and as the response for the color report control task (see experimental paradigm and timing). Participants wore headphones to hear auditory instructions and cues. During each trial, subjects were in complete darkness apart from the visual stimuli, which were not bright enough to illuminate the workspace. The hand was never visible to the subject, even during reaching.

There were 3 types of visual stimuli presented by different colors: the fixation point in yellow, targets in green or red, and masks in white. All stimuli were presented horizontally on the touchscreen and had the same diameter of 3 mm as the optic fibres. There was one central fixation location. Eight horizontal peripheral targets (4 on each side of the touchscreen) were used (Figure 1B), and twenty “mask” LEDs were located above and below the target line (ten on each side with five above and five below the targets). The visual mask was used during the delay periods to control for visual afterimages. The distance between the eyes of the subject and the center of the touchscreen was ∼60 cm. The target LEDs were located approximately 4°, 5°, 6° or 7° to the left or right of the fixation LED.

### Experimental paradigm and timing

We used an event-related design, with each trial lasting 38 seconds (including an inter-trial interval of 12 seconds). The paradigm included 3 tasks: reach, saccade, and color report as a control (Figure 1B). Each trial began with the presentation of the yellow fixation LED (this was displayed for 24 seconds before the first trial in each run). Concurrently, subjects were given the auditory instruction “move” or “color” to indicate the type of task they had to perform at the end of that trial. The important distinction between these two instructions is that while remembering the spatial location of the target LED (the visual target) was required for the movement trials, this information could be ignored for the color report trials. After 2 seconds, a green or red target LED was illuminated for 2 seconds, followed by an 8 second delay period (the effector-agnostic ‘visual target representation’ phase) during which the fixation LED and mask LEDs were illuminated. At the end of the delay, subjects were given one of 3 auditory instructions. For movement trials: “reach” (indicating a reach trial) or “look” (indicating a saccade trial). For color report trials the instruction “color” was repeated. This took 2 seconds. The independent effector instruction being given in the middle of the trial prevented subjects from forming their movement plan during the first delay period. The auditory instruction was followed by another 8 second delay period (the ‘movement planning’ phase) during which the fixation LED and mask LEDs were illuminated. After the mask LEDs were turned off, subjects heard a beep that served as a ‘go’ signal for subjects to move their arm or eye to the remembered location of the target in movement trials, or press the button once if the target LED was green or twice if it was red for the color report trials (or vice versa, this was be counterbalanced across subjects). This is referred to as the ‘movement execution’ phase. After touching the touchscreen or fixating the target location for 2 seconds, subjects heard a beep that instructed them to return their arm or eye to the starting position. The following trial started 12 seconds later.

Each functional run consisted of 12 trials presented in a random order (4 for each of the three tasks; 50% of targets presented in each visual hemifield for each task) and lasted about 8 minutes. For analysis, target locations were collapsed together as “left” or “right”. Subjects participated in 8 functional runs in one session. They were trained to perform the required tasks 1-2 days before imaging and practiced all tasks within the MRI scanner before scanning to ensure that they were comfortable with the task.

### Behavioral recordings

Following the fMRI experiments, the eye position and reach endpoints were inspected. Eye movement errors were defined as trials where subjects were unable to maintain visual fixation from target presentation until touching the touchscreen, or when the eye moved to the direction opposite of the instructed saccade goal. Reaching errors were defined as reaches to the direction opposite to the instructed reach goal.

### Imaging parameters

The experiment was conducted at the York MRI Facility at the Sherman Health Sciences Centre at York University with a 3-T whole-body MRI system (Siemens Magnetom TIM Trio). The posterior half of a 12-channel head coil (6 channels) was placed at the back of the head, with a 4-channel flex coil over the anterior part of the head (Figure 1B). The head was tilted ∼20° to allow for direct viewing of the stimuli during experimental trials.

Functional data was acquired using an EPI (echo-planar imaging) sequence (repetition time [TR] = 2000 ms; echo time [TE] = 30 ms; flip angle [FA] = 90°; field of view [FOV] = 192 mm × 192 mm, matrix size = 64 × 64 leading to an in-slice resolution of 3 mm × 3 mm; slice thickness = 3.5 mm, no gap; 36 transverse slices angled at ∼25° covering the whole brain). Slices were collected in ascending and interleaved order. During each experimental session, a T1–weighted anatomical reference volume was acquired using an MPRAGE sequence (TR = 1900 ms; TE = 2.52 ms; inversion time TI = 900 ms; FA = 90°; FOV = 256 mm × 256 mm × 192 mm, voxel size = 1 × 1 × 1 mm^3^).

### Preprocessing

All data was analyzed using BrainVoyager QX 2.2 (Brain Innovation). The first 2 volumes of each scan were discarded to avoid T1 saturation effects. For each run, slice scan time correction (cubic spline), temporal filtering (removing frequencies <2 cycles/run) and 3D motion correction (trilinear/sinc) were performed. The 3D motion correction was performed by aligning each volume of one run to the volume of the functional scan that was closest in time to the anatomical scan. 3 runs showing abrupt head movement of 1 mm or 1° were discarded. Functional runs were coregistered to the anatomical image. Functional data was then transformed into Talairach space using the spatial transformation parameters from each individual subject’s anatomical scan. The voxel size of the native functional images was 3×3×3 and was not resampled to a different voxel size during the preprocessing steps. Functional data was spatially smoothed using a FWHM of 8 mm.

### Data analysis

For each participant, we used a general linear model with 33 predictors. Two predictors were used for the initial auditory instruction (move or color); four predictors were used for visual target presentation (left or right X move or color trial); four predictors were used for visual target representation (left or right X move or color trial); three predictors were used for the 2^nd^auditory instruction (reach, saccade, or color trial); six predictors were used for motor preparation (left or right X reach, saccade, or color trial); six predictors were used for motor execution (left or right X reach, saccade, or color trial). In addition, six motion correction parameters and predictors for behavioral errors and inter-trial intervals were added as confound errors. Each predictor was derived from a rectangular wave function convolved with a standard hemodynamic response function using BrainVoyager QX’s default double-gamma hemodynamic response function.

### Voxelwise analysis

Contrasts were performed on β weights using an RFX (random effects) GLM with a percentage signal change transformation. This GLM was used to investigate the first two main questions for this study. To investigate the brain areas involved in effector-independent visual target representation and effector-dependent reach and saccade movement planning and movement execution, we performed five contrasts to find brain areas that showed higher activity for movement trials (reach and saccade) than the control (color) trials during each phase. We also performed two additional contrasts to test if brain areas showed effector-specific activation for reaches or saccades during movement planning and execution.

Activation maps for group voxelwise results were overlaid on the inflated brain of one representative subject. To correct for multiple comparisons, cluster threshold corrections (Forman et al., 1995) were performed for each contrast using BrainVoyager QX’s cluster-level statistical threshold estimator plug-in (1000 iterations). Areas that did not survive were excluded from further analysis. A Bonferroni correction was applied to the t value for each contrast to account for the two types of contrasts performed in the experiment: 1) movement trials > control trials (reach > control; saccade > control), and 2) effector specificity contrasts (reach > saccade). These two types of contrasts were planned a priori, with contrasts 1-5 being movement > control trials at three different time periods (1 effector independent and 2 effector-dependent) and contrasts 6 & 7 investigating effector-specificity during the planning and execution phases. (α = 0.05 / 2 comparisons = 0.025 corrected for *p* < 0.05).

## Results

To examine reach and saccade planning, we looked at general, non-directional movement activation; combining left and right movements for reach, saccade, and the colour control task. Figure 2 plots the effector-agnostic activation during the visual target representation phase, with the corresponding Talairach coordinates shown in Table 1. Figure 3 plots the independent effector activation for both reaches and saccades (vs the control task) during the planning and execution phases, with the corresponding Talairach coordinates shown in Tables 2 and 3. Brain areas were labeled by comparing the Talairach coordinates from the peak voxel within a cluster and comparing it to known sites of activation in the visuomotor system. It is important to note that at this point certain effector-specific overlapping functional areas have been labeled based on the effector used in the task (e.g., frontal eye fields vs. dorsal premotor cortex in frontal cortex). These data are described in more detail in the following sections. For a complete list of abbreviations for regions of interest (ROI) discussed in this study, see table 1.

**Table 1.**
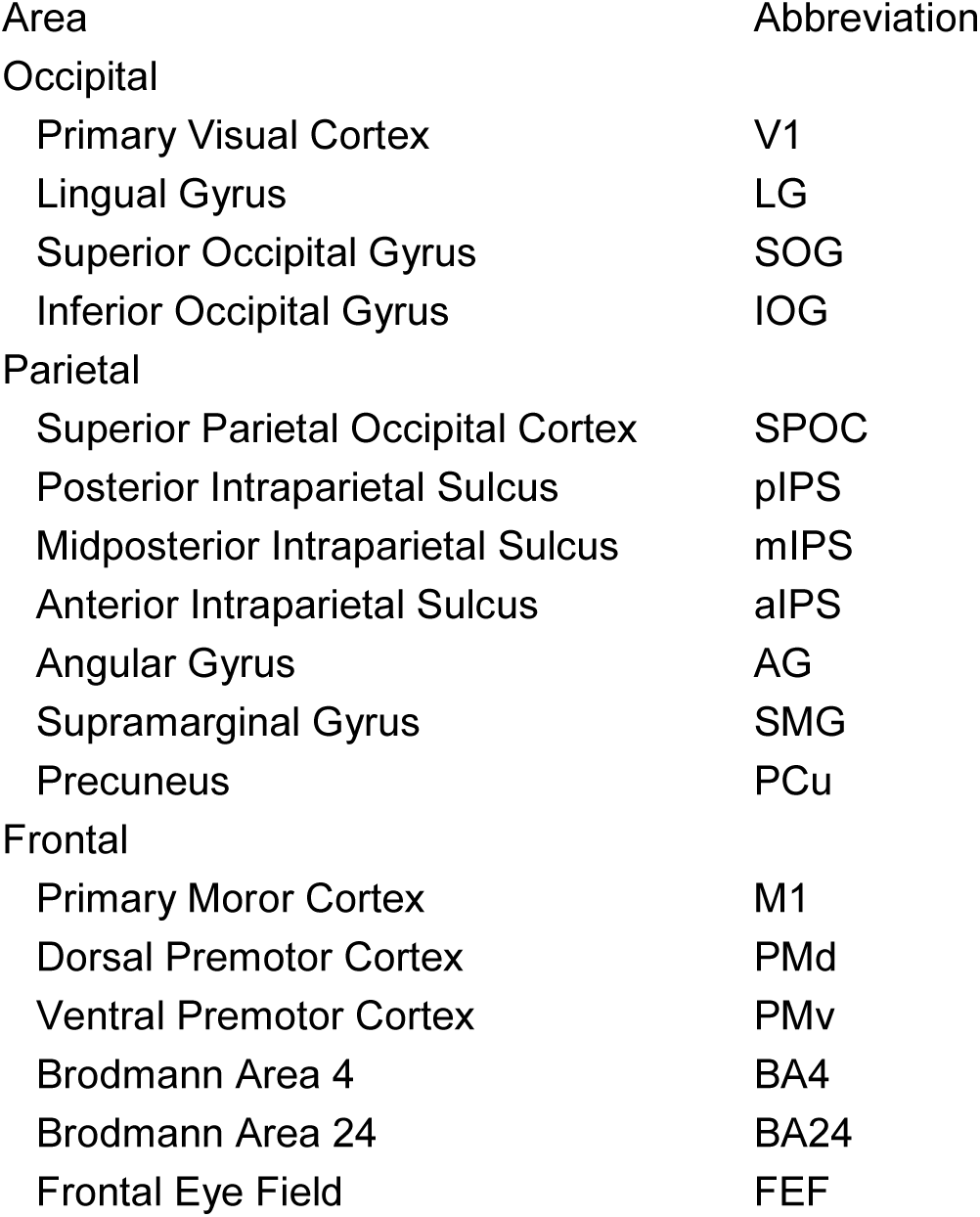
List of ROI brain area abbreviations

**Table 2.** Talairach coordinates and number of voxels for ROIs from the effector-agnostic target representation phase

| Area | Mean X | Mean Y | Mean Z | Voxels |
| --- | --- | --- | --- | --- |
| Left mIPS | -30.5 | -58.99 | 47.1 | 366 |
| Left AG | -49.76 | -47.12 | 12.59 | 464 |
| Right mIPS | 20.82 | -61.59 | 50.39 | 617 |

**Table 3.** Talairach coordinates and number of voxels for ROIs from the independent effector movement planning phase

| Area | Mean X | Mean Y | Mean Z | Voxels |
| --- | --- | --- | --- | --- |
| Reach Areas |  |  |  |  |
| Left PMd | -26.73 | -6.3 | 57.55 | 422 |
| Left M1 | -23.2 | -29.79 | 63.76 | 370 |
| Left BA4 | -7.03 | -22.43 | 44.92 | 672 |
| Left aIPS | -13.84 | -43.84 | 61.19 | 342 |
| Left mIPS | -18.85 | -48.09 | 52.67 | 688 |
| Left pIPS | -23.33 | -73.21 | 44.08 | 350 |
| Left SPOC | -14.45 | -72.06 | 41.09 | 519 |
| Left SOG | -20.43 | -78.7 | 26.72 | 846 |
| Left MOG | -33.25 | -70.6 | 10.3 | 443 |
| Left IOG | -41.42 | -67.91 | -4.36 | 237 |
| Right PMd | 21.76 | -11.53 | 58.03 | 710 |
| Right mIPS | 21.44 | -57.75 | 52.86 | 338 |
| Right pIPS | 12.3 | -72.25 | 46.49 | 257 |
| Saccade Areas |  |  |  |  |
| Left pIPS | -17.04 | -69.09 | 48.84 | 616 |
| Left SOG | -24.67 | -84.38 | 19.38 | 355 |
| Left MOG | -32.89 | -81.34 | 3.02 | 221 |
| Right pIPS | 21.44 | -68.48 | 36.94 | 441 |
| Right MOG | 31.1 | -76.62 | 9.19 | 294 |
| Right SOG | 21.36 | -81.64 | 22.22 | 318 |

**Table 4.**
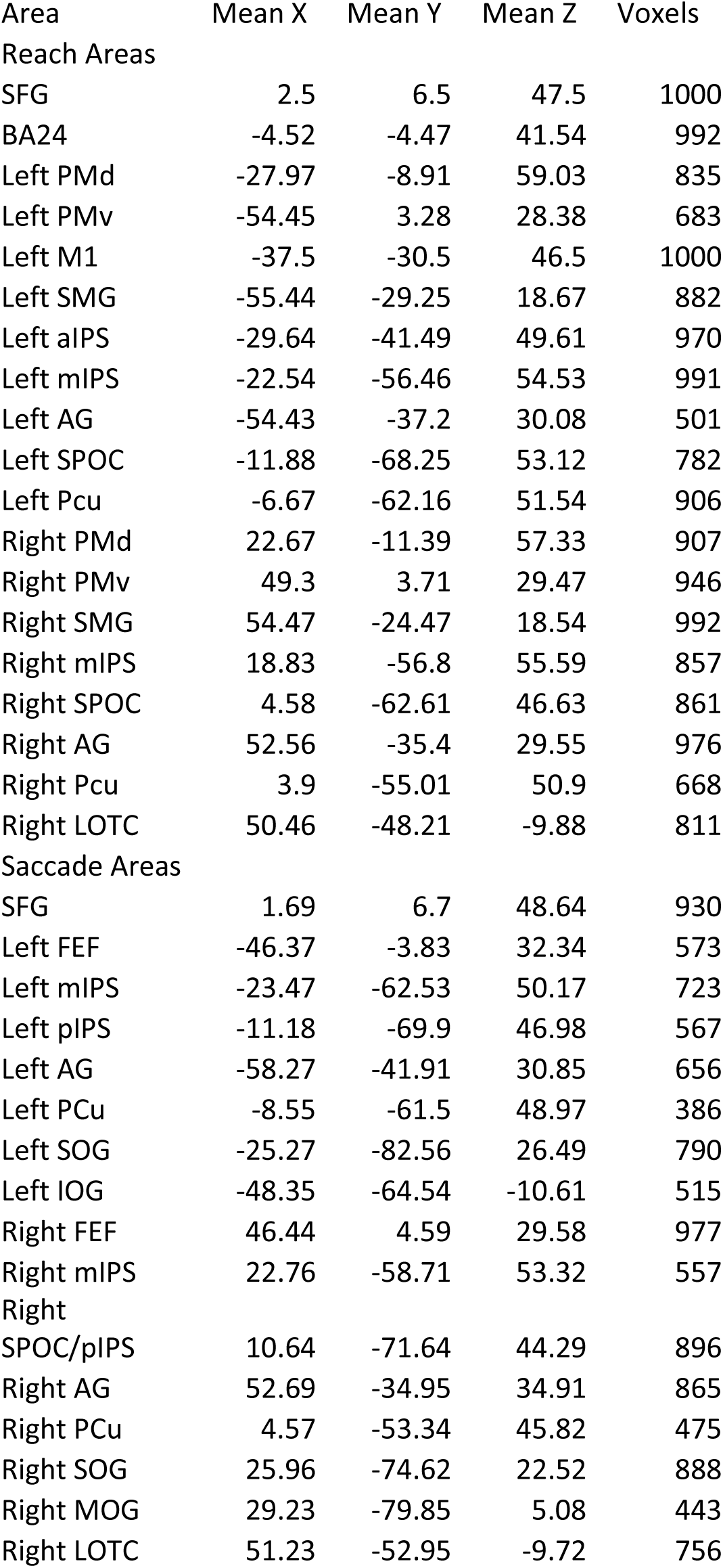
Talairach coordinates and number of voxels for ROIs from the independent effector movement execution phase

**Figure 2.**
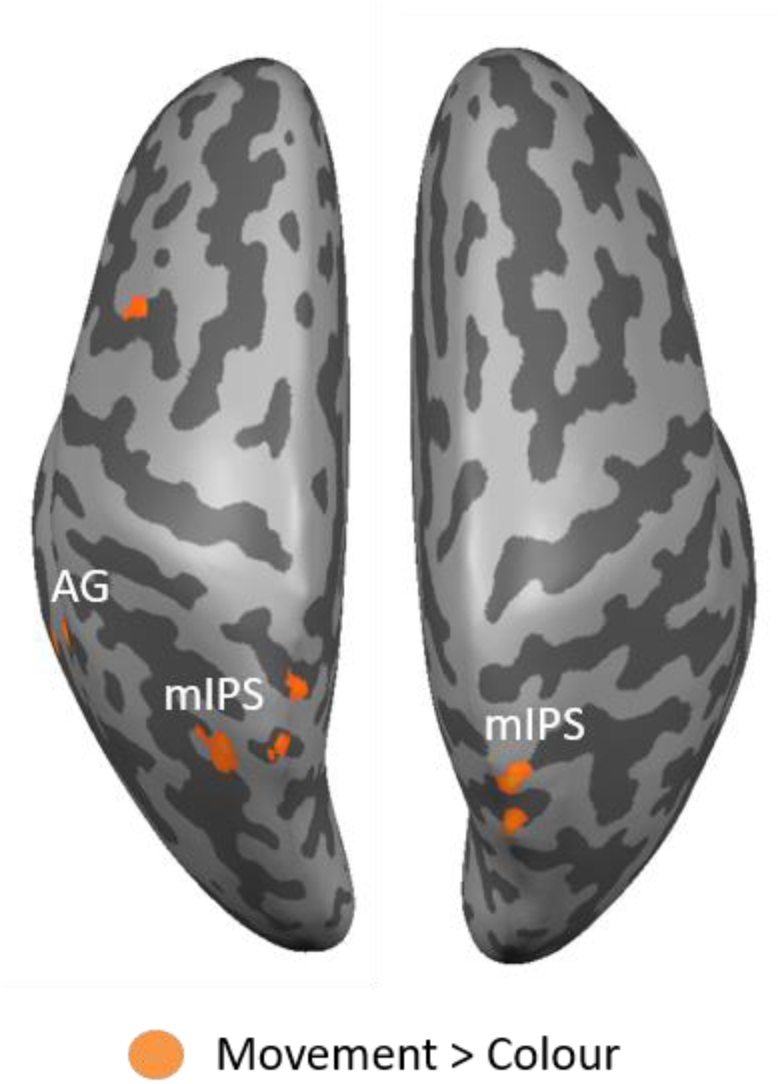
Effector-agnostic target representation phase (before the effector is known). Voxelwise statistical maps obtained from the RFX GLM for the contrast Movement (Reach+Saccade) > Color report. Event-related group activation maps for target representation are displayed on the ‘inflated brain’ of one representative subject, where light gray represents gyri and dark gray represents sulci. The leftward inflated brain represents the left hemisphere, and the rightward brain represents the right hemisphere. Highlighted areas show significantly higher activation than control data with a p<0.05 with Bonferroni and cluster threshold corrections. These areas include the left and right mIPS and Left AG.

**Figure 3.**
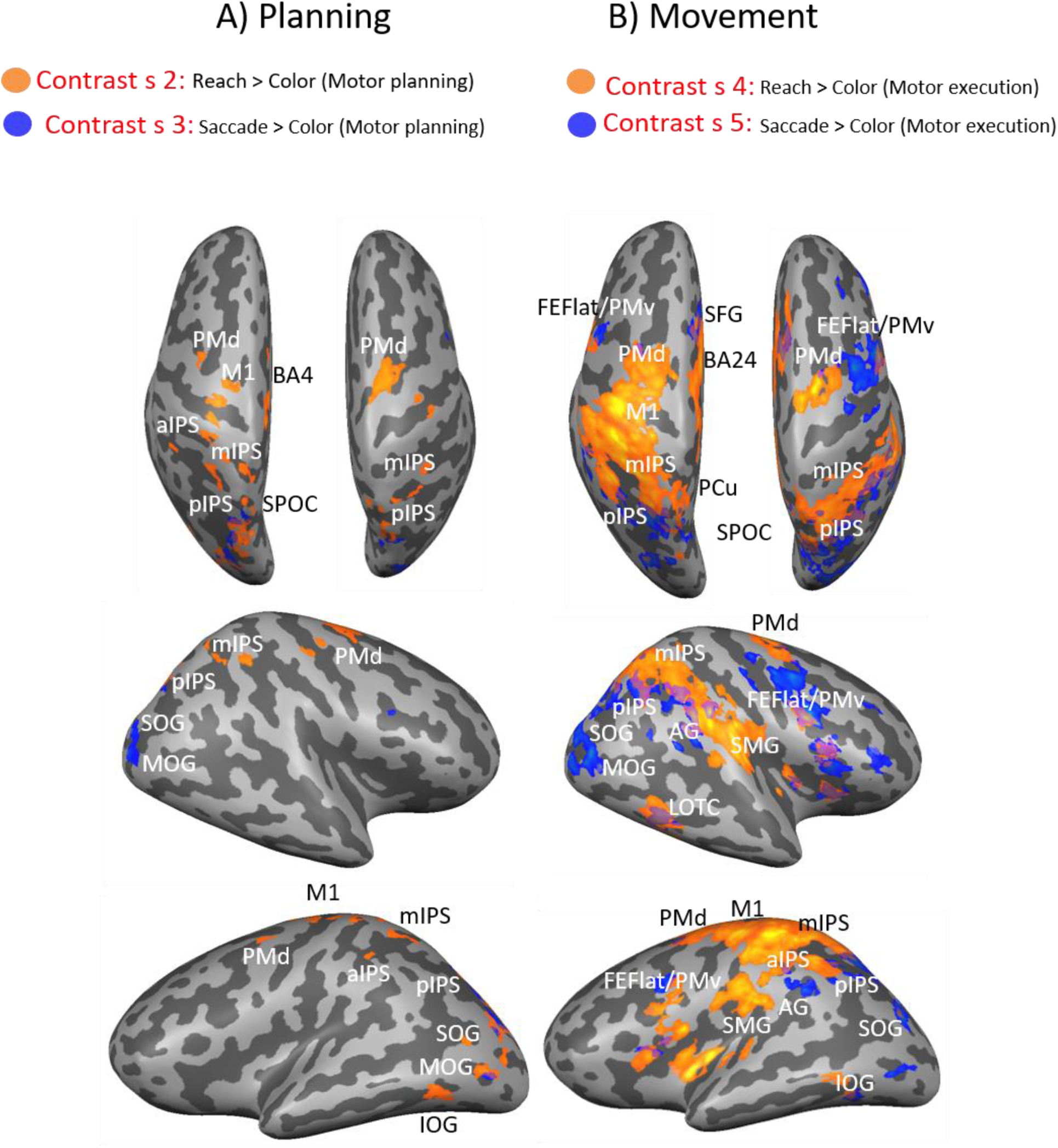
Independent Effector movement planning and execution (once the effector is known). A) Voxelwise statistical maps obtained from the RFX GLM for the contrasts Reach > Color report (orange) and Saccade > Colour report (blue). Event-related group activation maps are displayed on the inflated brain of one representative subject for movement planning. Highlighted areas show significantly higher activation than control data with a p<0.05 with Bonferroni and cluster threshold corrections. These areas include for reach bilateral PMd, mIPS, pIPS, and SOG. Significant activation was also observed in left M1, SPOC, and IOG. For Saccades, significant activation was observed in bilateral pIPS and SOG. B) Voxelwise statistical maps obtained from the RFX GLM for the contrasts Reach > Color report (orange) and Saccade > Colour report (blue). Event-related group activation maps are displayed on the inflated brain of one representative subject for movement execution. Highlighted areas show significantly higher activation than control data with a p<0.05 with Bonferroni and cluster threshold corrections. These areas include for reach bilateral PMd, PMv, mIPS, SMG, IOG, SMA and IFG. Significant activation was also observed in left M1. For Saccades, activation was observed in bilateral FEF, mIPS, pIPS, SPOC, AG, and SOG. (See Table 1 for site abbreviations)

### Effector-agnostic activation during the Target representation phase

Note that in our paradigm subjects could not predict which effector they would use in the first Target Representation delay, so one might expect activation related to target memory, general motor preparation, or motor preparation required for both effectors. Contrast 1 [Target Representation Move > Target Representation Color] investigated which brain areas showed higher activation for visuospatial coding required to plan a target-directed movement (either reach or saccade) than activation related to representing the color of the target (the requirement of the control task). In this phase, only the target location was known (as the effector was only specified by an auditory instruction after this delay period), and any activation revealed by this contrast may be related to any aspect of target coding (not limited to spatial location) or early effector-independent movement preparation. Figure 2 shows the activation map for this contrast superimposed on inflated cortical surfaces viewed from above. The indicated areas survived a cluster threshold correction of 15 voxels. This contrast revealed modest bilateral activation near the midposterior intraparietal sulcus (mIPS), and modest unilateral activation in the left angular gyrus (AG). At first glance it might seem odd that only areas associated with movement control (Gallivan and Culham, 2015) were activated, but recall that the control task also involves memory of a non-spatial, non-motor target type. Thus, this subtraction shows areas with memory-epoch activity *specific to spatial location or early effector-independent motor preparation for saccade, reach, or both*.

### Independent Effector activation versus Color Control During Planning and Execution

Following the target representation phase, subjects were provided with an effector cue (Figure 1), allowing us to test the influence of this cue on BOLD activation during motor planning. To do this, we started with two independent, independent effector contrasts: <u>Movement planning phase</u>: Contrast 2 [Movement Planning Reach > Movement Planning Color] and Contrast 3 [Movement Planning Saccade > Movement Planning Color] investigated which brain areas showed higher activation for movement planning for reach and saccade, respectively, than activation related to representing the color of the target (the requirement of the control task). Activation during this phase could be related to planning a specific movement and/or general motor preparation in anticipation of an upcoming reach (for Contrast 2) or saccade (for Contrast 3). The activation map for these contrasts are shown on an inflated cortical surface viewed from above and the lateral view (Figure 3a), with orange activation representing reach activation and blue activation representing saccade activation. The marked areas survived a cluster threshold correction of 26 voxels for Contrast 2 (reach), and 27 voxels for Contrast 3 (saccade). For reach, Contrast 2 revealed modest bilateral activation near the intersection of the precentral and superior frontal sulci, consistent with the location of dorsal premotor cortex (PMd) (Monaco et al. 2011), midposterior intraparietal sulcus (mIPS), and posterior intraparietal sulcus (pIPS). Activation was also found in the left hemisphere in primary motor cortex (M1), Brodmann Area 4 (BA4), superior parietal occipital cortex (SPOC), superior occipital gyrus (SOG), middle occipital gyrus (MOG), and inferior occipital gyrus (IOG). For a complete list of abbreviations for regions of interest (ROI) discussed in this study, see table 1. For Saccade, Contrast 3 revealed modest bilateral activation for posterior intraparietal sulcus (pIPS) and superior occipital gyrus (SOG). A conjunction analysis was also performed to investigate any ROIs involved in both processes, however no regions were found to be significantly active for both effectors.

#### Movement execution phase

Contrast 4 [Movement Execution Reach > Movement Execution Color] and Contrast 5 [Movement Execution Saccade > Movement Execution Color] investigated which brain areas showed higher activation related to executing a reach or saccade, respectively, than activation related to indicating the color of the target with a button press (the requirement of the control task). The activation map for these contrasts are shown on an inflated cortical surface (Figure 3B). The marked areas survived a cluster threshold correction of 43 voxels for Contrast 4 (reach), and 29 voxels for Contrast 5 (saccade). For reach, Contrast 4 revealed widespread activation in bilateral mIPS, pIPS, SPOC, AG, PMd, PMv, Brodmann Area 24 (BA24), superior frontal gyrus (SFG), supramarginal gyrus (SMG), and AG. Activation was also found in the left hemisphere in precuneus (PCu), aIPS, M1, SOG, and IOG, and in the right hemisphere in lateral occipital temporal cortex (LOTC). For saccades, Contrast 5 revealed activation in bilateral lateral frontal eye fields (FEF), mIPS, pIPS, SPOC, AG, and SOG, as well as right LOTC. A conjunction analysis on contrasts 4 and 5 was done to investigate any ROIs involved in both processes, and the results can be found in Figure 4. Left mIPS and right AG, PCu, SPOC, LOTC, and FEF/PMv were found to be significantly active for both effectors.

**Figure 4.**
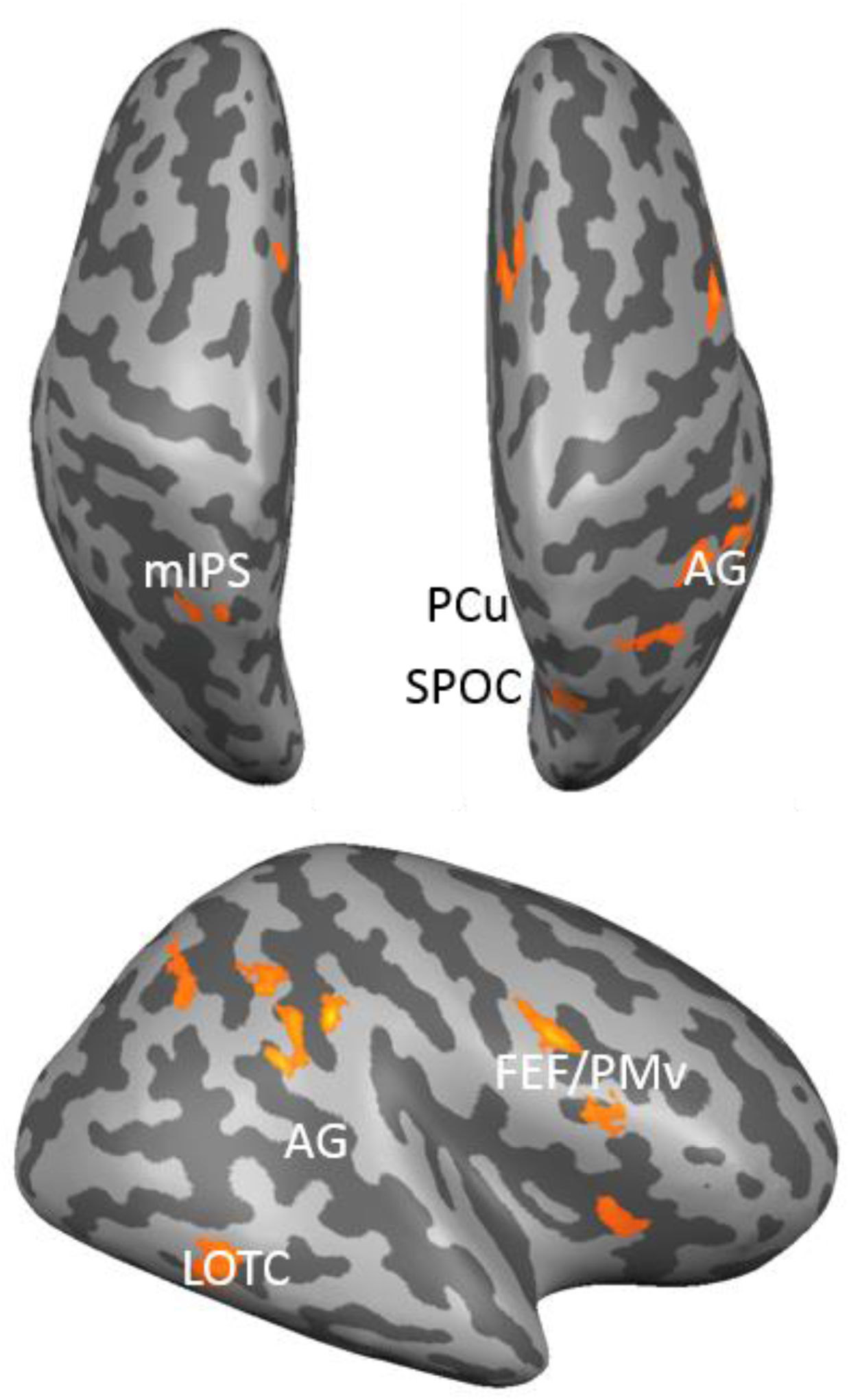
A conjunction analysis on contrasts 4 and 5 (reach>colour and saccade>colour during movement execution. Event-related group activation maps are displayed on the inflated brain of one representative subject for movement planning. Highlighted areas show significantly higher activation than control data with a p<0.05 with Bonferroni and cluster threshold corrections. Left mIPS and right AG, PCu, SPOC, LOTC, and FEF/PMv were found to be significantly active for both effectors.

### Independent Effector Time Series Analysis

To provide a more detailed understanding of the temporal progression of BOLD signals during our task, we examined time series data derived from the saccade and reach regions of interest identified in the movement execution phases in Figure 3.

Figure 5 illustrates the time course data of the reach and color conditions for 12 brain areas from Contrast 4, chosen because they have been linked to visuomotor reach planning, including: left and right mIPS, left and right SPOC, left and right AG, left and right SMG, left and right PMd, left M1, and left SOG. Figure 6 illustrates the time courses data of the saccade and color conditions for 10 brain areas from Contrast 5, chosen because they have been linked to visuomotor saccade planning, including: left and right FEF, left and right mIPS, left and right SPOC, left and right AG, and left and right SOG.

**Figure 5.**
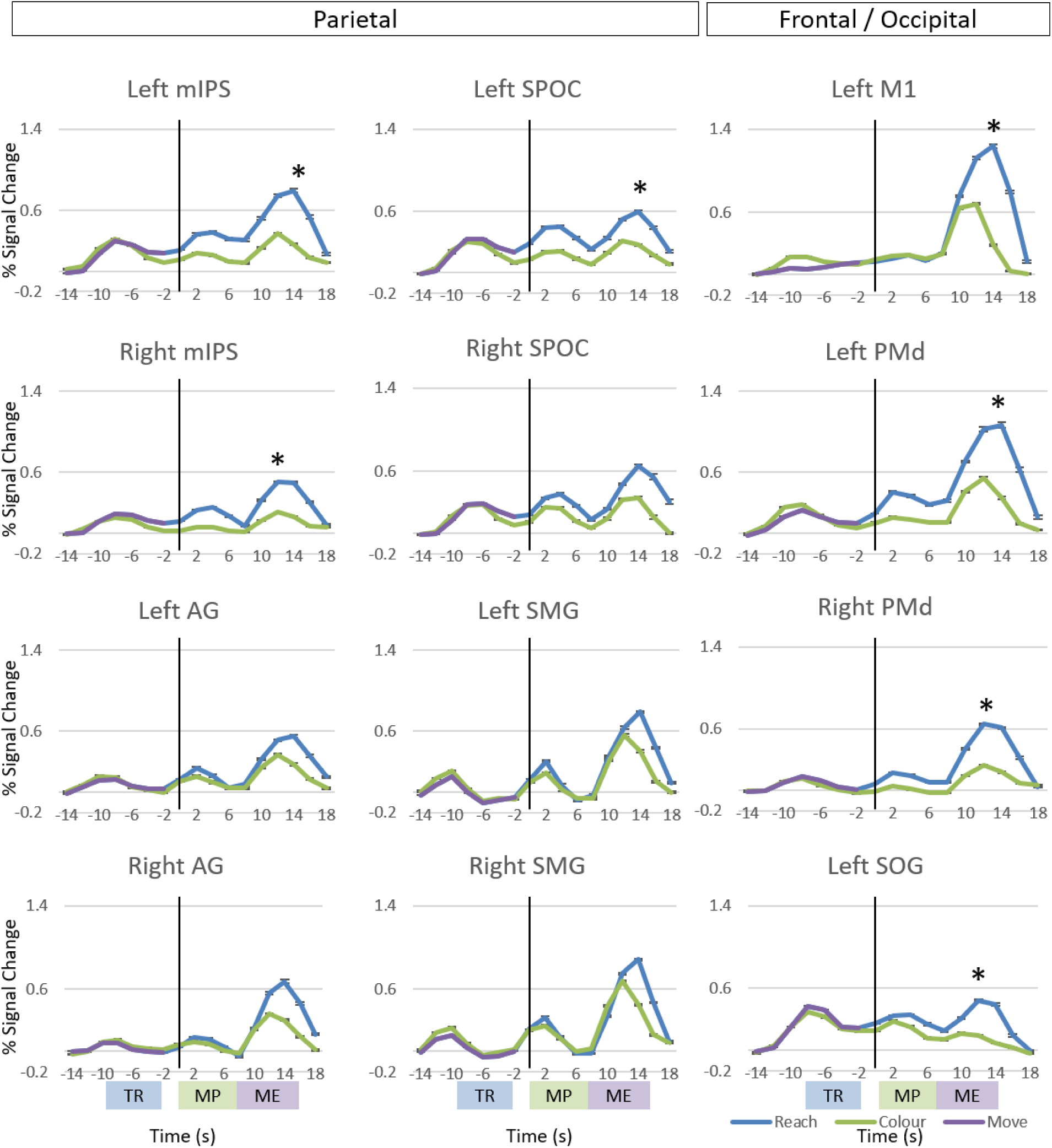
Time courses for brain areas of interest (bilateral mIPS, SPOC, AG, SMG, and PMd; left M1 and SOG) that were active from the Reach > Color contrast during the movement execution phase. The blue line indicates activity (% signal change) from reach trials and the green line indicates activity from color report trials. The purple line indicates activation in reach trials before the effector was known (general pre-movement activation). Error bars are SEM across subjects. The x axis displays time in seconds and is time locked to the movement planning phase. The vertical black line indicates the onset of the movement planning (MP) phase, while the Target Representation (TR), movement planning, and Movement Execution (ME) phases are identified along the x axis (from left to right). An asterisk (*) indicates a significant difference between the reach and colour activation at the time peak reach activation.

**Figure 6.**
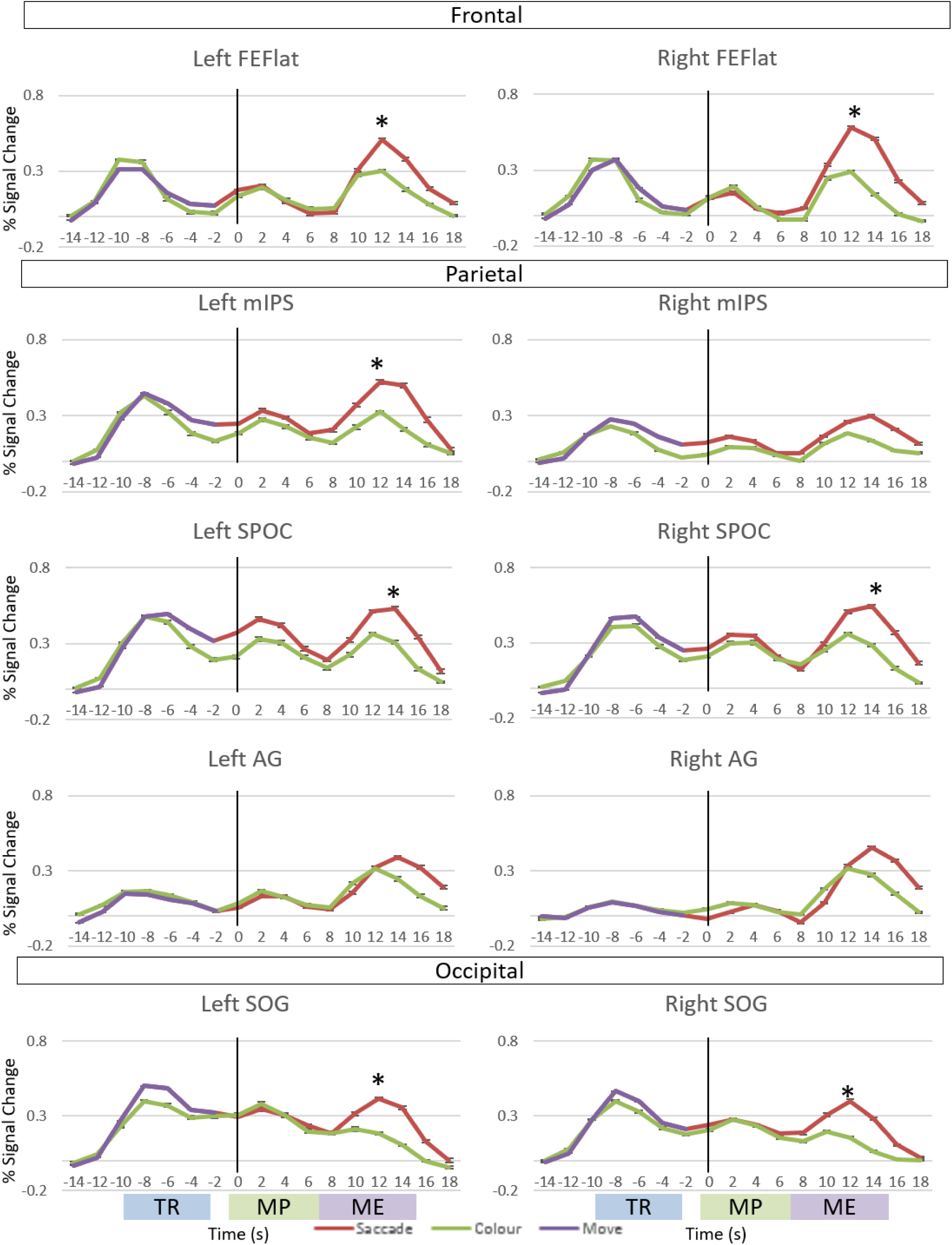
Time courses for brain areas of interest (bilateral FEF, mIPS, SPOC, AG, and SOG) that were active from the Saccade > Color contrast during the movement execution phase. The red line indicates activity (% signal change) from saccade trials and the green line indicates activity from color report trials. The purple line indicates activation in reach trials before the effector was known (general pre-movement activation). Error bars are SEM across subjects. The x axis displays time in seconds and is time locked to the movement planning phase. The vertical black line indicates the onset of the movement planning (MP) phase, while the Target Representation (TR), movement planning, and Movement Execution (ME) phases are identified along the x axis (from left to right). An asterisk (*) indicates a significant difference between the reach and colour activation at the time peak saccade activation.

Looking at these time courses, several patterns emerge that help to understand the previous observations and provide reference events for further analysis. First, all regions of interests show three peaks of activation, aligned closely with target representation, movement planning, and movement execution phases. Second, the relative heights of these peaks were dependent on the known functional role of each area, with SOG (in occipital cortex) showing a relatively larger target peak (than ‘planning’ and ‘execution’), SPOC showing roughly equal target, planning, and execution peaks, and mIPS, PMd, AG, FEF, and M1 showing predominant movement execution peaks. Third, the degree of movement task-specificity (gap between task vs. control lines) generally increased both in time from visual target representation to movement execution and in cortical space from occipital cortex to parietal cortex to frontal cortex. Thus, the entire occipital-parietal-frontal axis was activated during target coding, planning, and execution, but the task-specificity of these responses increased along the antero-frontal axis and in the temporal transition from target, planning, and execution responses.

To quantify independent effector activation in the time courses in figures 5 and 6, we performed paired two tailed t-tests between the reach and colour (figure 5) and saccade and colour (figure 6) data at the time of peak independent effector activation (the maximum % signal change value once the effector has been specified for the trial) to indicate significant reach or saccade activation, respectively. We limited our comparisons to this time point to indicate the presence of independent effector activation without needing to correct for multiple comparisons across all time points.

As one might predict, the peak activation for all reach and saccade areas following the specification of the effector to be used in the trial occurred during motor execution. For reach (figure 5), the peak activation during reach execution was significantly higher than the colour activation at the same time point in bilateral mIPS, bilateral PMd, left SPOC, left M1, and left SOG. For saccades (figure 6), the peak activation during saccade execution was significantly higher than the colour activation at the same time point in bilateral FEF, bilateral SPOC, bilateral SOG, and left mIPS. For both reach and saccade, we observed significant differences between the experimental and control conditions during motor execution in areas that have been linked to reach and saccade planning. It is worth noting, however, that the overall shapes of the curves for all these brain areas were similar (including those that did not reach significance), indicating a degree of planning and execution activation across frontal, parietal, and occipital areas for both reach and saccade. To better understand how this activation differs for reaches and saccades, the next section examines effector-specific activation during planning and execution.

### Effector-Preference Direct Contrasts: Reach vs. Saccade Activation for Planning and Execution

While our first set of contrasts on the movement planning and execution phases assessed independent effector activation, we also performed effector-preference contrasts to directly compare reach and saccade activation. Figure 7 plots the effector-preference activation during the planning and execution phases.

**Figure 7.**
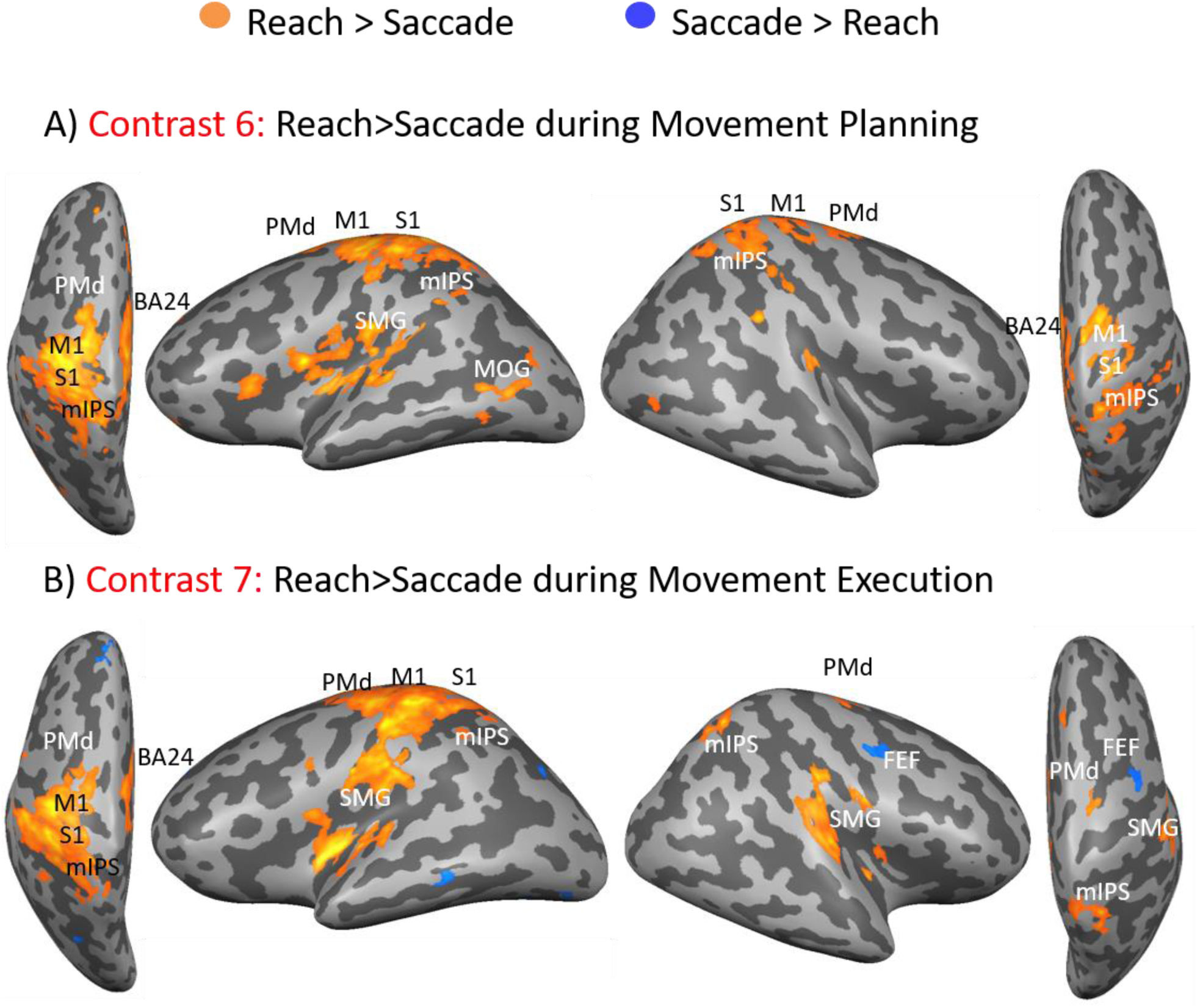
Effector-preference movement planning and execution. A) Voxelwise statistical maps obtained from the RFX GLM for the contrasts Reach > Saccade. Event-related group activation maps are displayed on the inflated brain of one representative subject for movement planning. Highlighted areas show significantly higher reach activation than saccade data with a p<0.05 with Bonferroni and cluster threshold corrections. These areas include for reach bilateral PMd, mIPS, S1, and BA24, as well as left MOG and SMG. B) Voxelwise statistical maps obtained from the RFX GLM for the contrasts Reach > Saccade. Event-related group activation maps are displayed on the inflated brain of one representative subject for movement execution. Highlighted areas show significantly higher activation than control data with a p<0.05 with Bonferroni and cluster threshold corrections. These areas include for reach bilateral PMd, mIPS and SMG, as well as left BA24, M1, and S1. For Saccades, significantly higher activation was observed in right FEF.

#### Effector-preference activation during the movement planning phase

Contrast 6 [Movement Planning Reach > Movement Planning Saccade] investigated which brain areas showed higher activation for movement planning for reaches than for saccades. The activation map for this contrast is shown on an inflated cortical surface viewed from above and the lateral view (Figure 7a), with orange activation representing reach activation and blue activation representing saccade activation. The marked areas survived a cluster threshold correction of 32 voxels. During movement planning, no saccade areas survived cluster threshold corrections. Several reach-specific areas were identified, including bilateral mIPS, S1, M1, PMd, and BA24, as well as left SMG and MOG.

#### Effector-preference activation during the movement execution phase

Contrast 7 [Movement Execution Reach > Movement Execution Saccade] investigated which brain areas showed higher activation for movement execution for reaches than for saccades. The activation map for this contrast is shown on an inflated cortical surface viewed from above and the lateral view (Figure 7b), with orange activation representing reach activation and blue activation representing saccade activation. The marked areas survived a cluster threshold correction of 38 voxels. Several reach-specific areas were identified, including bilateral mIPS and SMG, as well as left PMd, M1, S1, and BA24. The only saccade-specific area found was right FEF.

#### Time series data

To better understand the evolution of activation for these effector-preference brain areas at the time of movement execution, we examined their time series. We selected 8 ROIs from Contrast 7, including left and right PMd, left and right mIPS, left and right SMG, left M1, and left FEF (the only saccade-preference area). To quantify effector-preference activation differences in the time courses, we performed two tailed paired t-tests between the reach and saccade data at the time of peak effector-specific activation (the maximum % signal change value once the effector has been specified for the trial) to indicate significant reach-preference or saccade-preference activation. We limited our comparisons to this time point to indicate the presence of effector-preference activation without needing to correct for multiple comparisons across all time points. Looking at these time courses, one can start to see separation between saccade and reach planning during the movement planning phase. For the reach-specific areas, the increased activation observed in reach trials over saccade trials becomes quite pronounced during the motor execution phase, with all 7 areas showing significantly higher activation for reach than saccade trials. For right FEF, the activation appeared to be similar with the peak saccade activation not being significantly different than the reach activation at the same time point.

### Temporal correlation of effector preference between cortical areas

To quantify some of the qualitative observations made above, we performed temporal correlations of reach activation, saccade activation, and reach-saccade effector specificity activation between the regions identified in the movement execution effector-preference contrasts (from Figure 3B), which can be found in Figure 8. To do this, we used the % BOLD signal change time series data from the time when the effector-specific command is given (time 0) to 14 seconds after (to be inclusive of movement planning and movement execution phases). We then correlated between sites (r) by matching their BOLD signal changes for each scan in this time range.

**Figure 8.**
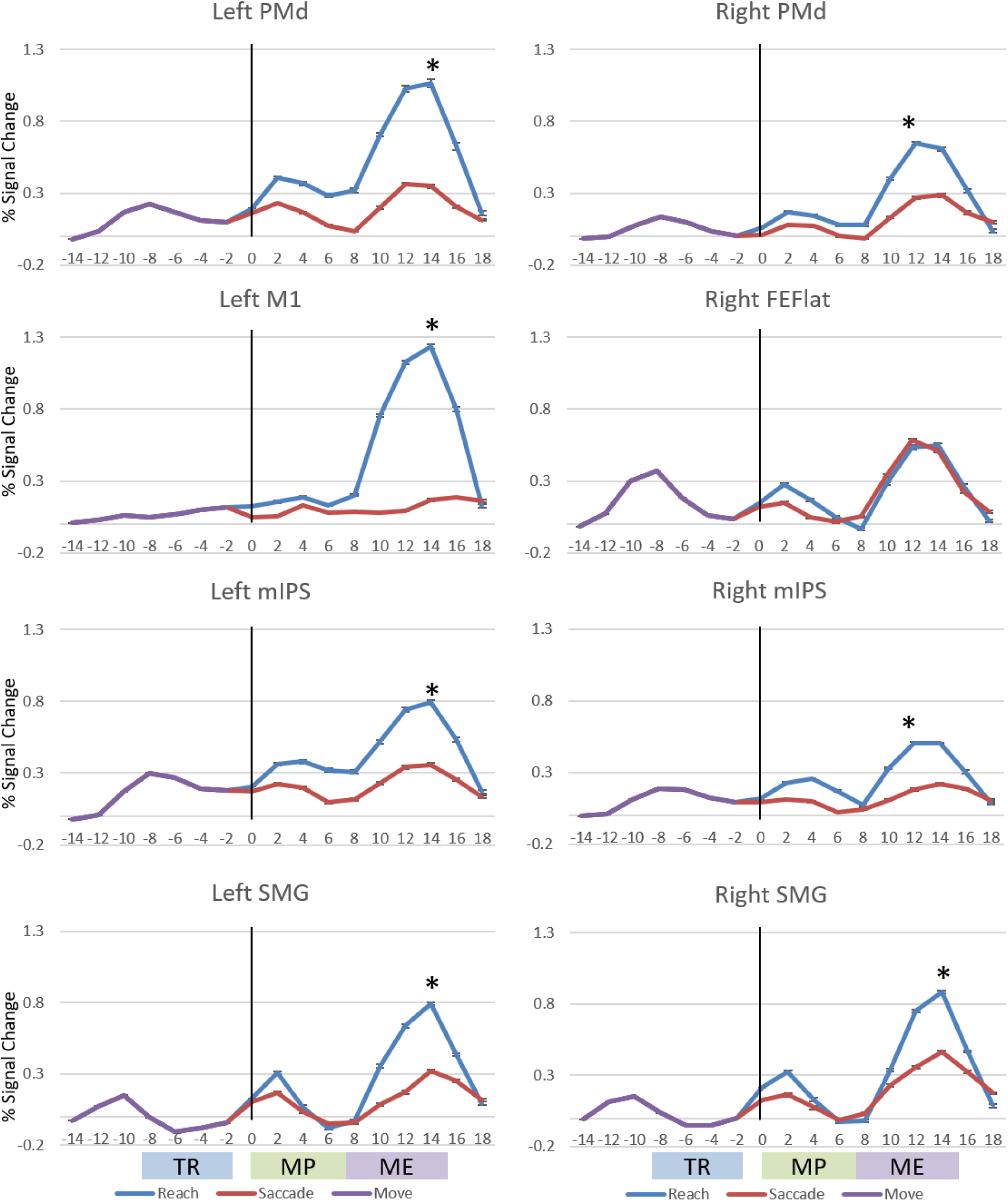
Time courses for brain areas of interest (bilateral PMd, mIPS, SMG; left M1; right FEF) that were active from the Reach > Saccade contrast during the movement execution phase. The blue line indicates activity (% signal change) from reach trials and the red line indicates activity from saccade trials. Error bars are SEM across subjects. The x axis displays time in seconds and is time locked to the movement planning phase. The vertical black line indicates the onset of the movement planning (MP) phase, while the Target Representation (TR), movement planning, and Movement Execution (ME) phases are identified along the x axis (from left to right). An asterisk (*) indicates a significant difference between the reach and colour activation at the time peak reach activation.

Given the hemispheric differences in reach planning found in our previous study (Cappadocia et al., 2017), we performed separate analyses for the left and right hemispheres.

Figure 9A shows the *saccade* effector preference correlations between each brain area for the right hemisphere and the left hemisphere. The brain areas have been ordered from posterior to anterior, and the variable shown is the r score. The resulting correlation matrix shows high r scores across the board, with notably high scores (indicated by a darker shade of red) for parietal brain regions (mIPS, PCu, AG). This was especially the case for mIPS, which consistently showed high (r > 0.9) correlations with all other saccade areas. These correlations were often significant (as indicated by bolded numbers) with a *p* < 0.05 with Bonferroni corrections for 6 comparisons [α = 0.05 / 6-1 comparisons = 0.01 corrected for *p* < 0.05].

**Figure 9.**
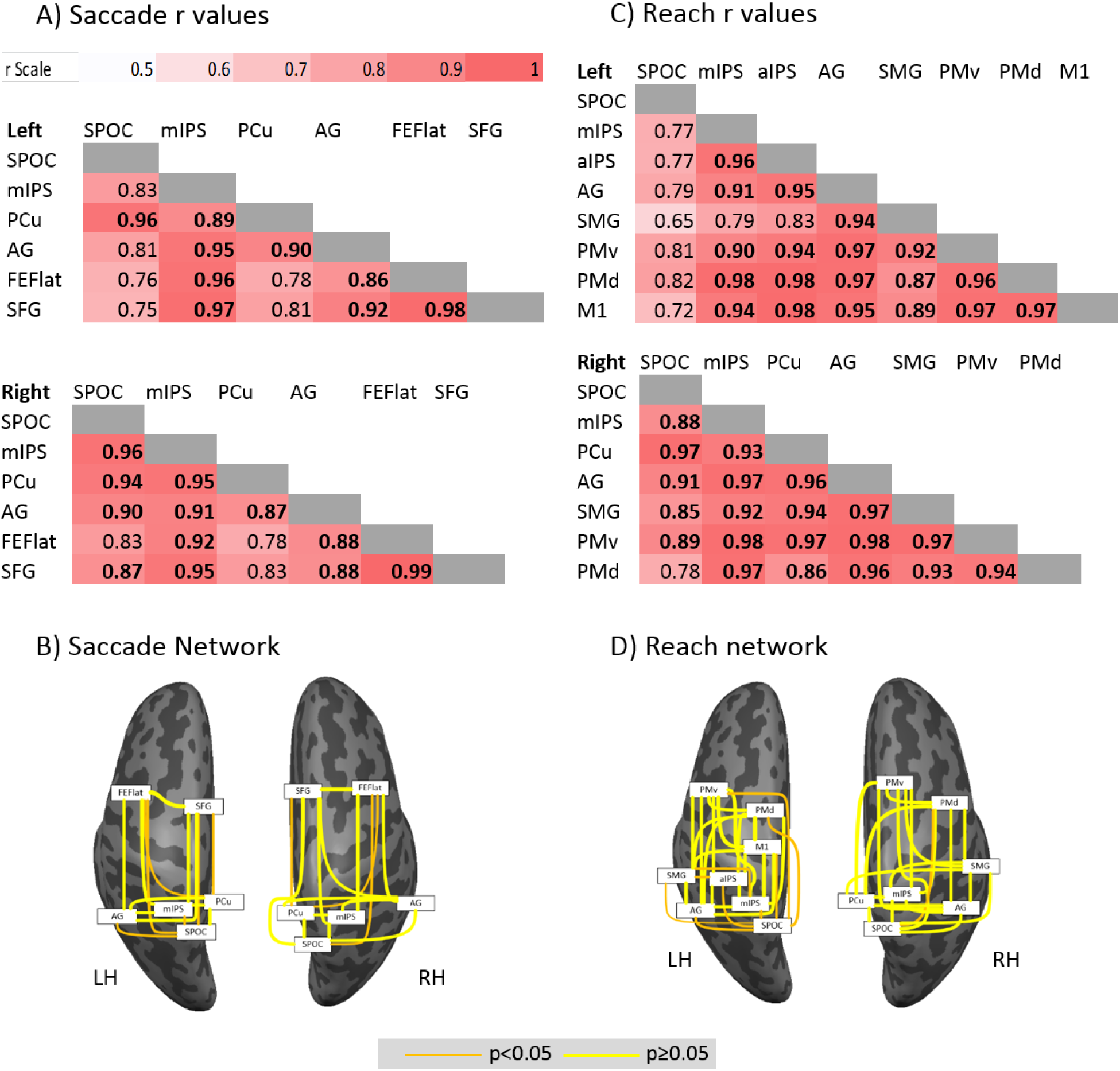
Correlations through time between regions derived from the saccade > colour report movement execution contrast. A) SACCADE: Matrices showing correlations (r) through time between all areas for each spatial domain tested (redundant entries in upper-right half are omitted) within each hemisphere. A continuous color scale is used to indicate the strength of correlation (r), i.e., with white close to 0.5 and red close to 1.0, and significant correlations (p<0.05, Bonferroni corrected) are bolded. B) Graphical representations of the strength of correlation between left and right hemisphere brain areas for saccades. The thickness of the line indicates the r^2^value, with a thin line being close to 0 and a thick line close to 1. For these plots we used r^2^to increase the difference between highly correlated and less correlated areas. These data are superimposed on left and right hemisphere ‘inflated brains’ from a typical subject where light gray signifies gyri and dark gray signifies sulci. C) REACH: Matrices showing correlations (r) through time between all areas for each spatial domain tested (redundant entries in upper-right half are omitted) within each hemisphere (same conventions as A). D) Graphical representations of the strength of correlation between left and right hemisphere brain areas for reach (same conventions as B).

Figure 9B graphically represents the same data as Figure 9A as a ‘network’ of correlations between our various regions of interest. The width of each line is scaled by the r^2^value for the two regions that it joins, with significant correlations highlighted in yellow (p < 0.05), non-significant correlations are shown in orange (p≥0.05). This figure also helps to visualize ‘hub’ areas in the visual domain, sprouting thick yellow lines (high correlations with yellow indicating significant correlations) toward numerous other areas, as opposed to thin orange lines (low correlations with orange indicating non-significant correlations). For saccades, one observes an extensive network of significant correlations in both hemispheres. Of note, FEF has non-significant correlations in both the left and right hemispheres to PCu and SPOC, and mIPS and AG appear as key hubs, with significant correlations to almost all regions.

Figure 9C similarly shows the effector preference *reach* correlations between each brain area. Again, areas were ordered from posterior to anterior. Most of our reach areas showed high (r > 0.9) and statistically significant correlations with all of the other areas (the exception being SMG, which was not significantly correlated with aIPS or mIPS). Another notable exception was left SPOC, which did not have significant correlations with any other areas in the left hemisphere. Similarly, in the right hemisphere, almost all areas had high and significantly correlated activation, with the one exception being the pairing of PMd and SPOC. These highly correlated networks can be viewed graphically in figure 9D, which demonstrates a much more extensive and densely correlated ‘network’ compared to the saccade ‘network’ illustrated in figure 8B.

Figure 10A shows the correlations *betwee*n reach and saccade ROIs within each hemisphere. This analysis was performed to assess the similarity in activation across effectors in each hemisphere. Again, areas were ordered from posterior to anterior, this time with saccade regions listed horizontally and reach regions listed vertically. Even at first glance, once can observe a stark difference between the hemispheres, where almost all right hemisphere brain areas (37/42) show statistically significant correlations, compared to the left hemisphere where only 15/48 regions (31%) are significantly correlated. In the left hemisphere, AG appears as a clear hub for both effectors, showing significant correlations with 7/8 reach areas and 3/6 saccade areas. Notable, left FEF did not have any significant correlations with any of the reach areas. These highly correlated networks can be viewed graphically in figure 10B, using the same methodology used for 17B.

**Figure 10.**
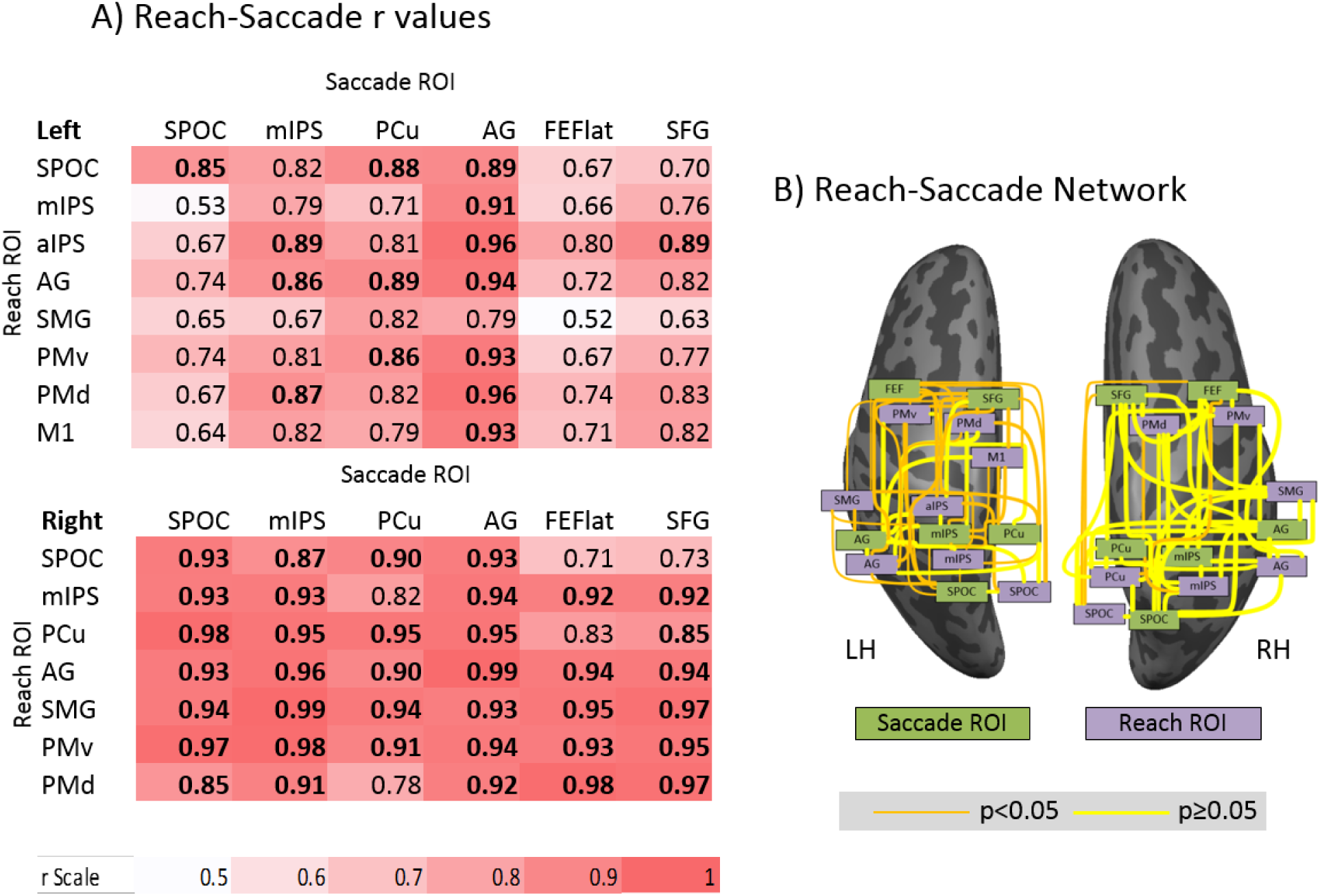
Correlations through time between regions derived from comparing the saccade > colour report ROIs to reach > colour report ROIs during the movement execution contrast. A) Matrices showing correlations (r) through time between all areas for each spatial domain tested within each hemisphere. A continuous color scale is used to indicate the strength of correlation (r), i.e., with white close to 0.5 and red close to 1.0, and significant correlations (p<0.05, Bonferroni corrected) are bolded. Reach areas are listed vertically and saccade areas are listed horizontally. B) Graphical representations of the strength of correlation between left and right hemisphere brain areas. The thickness of the line indicates the r^2^value, with a thin line being close to 0 and a thick line close to 1. For these plots we used r^2^to increase the difference between highly correlated and less correlated areas. These data are superimposed on left and right hemisphere ‘inflated brains’ from a typical subject where light gray signifies gyri and dark gray signifies sulci.

## Discussion

In this study, we used an event-related fMRI design to investigate several key questions. To summarize, the first was to identify which regions are differentially activated for effector-agnostic visual target representation, as well as independent effector reach and saccade movement planning and movement execution. This analysis revealed selective, bilateral mIPS and left AG activation during the effector-independent target representation phase. In the independent effector planning phase, the entire occipital-parietal-frontal reaching network was activated, while only bilateral pIPS and SOG activation was observed for saccades. For the independent effector movement execution phase, broad activation of occipital-parietal-frontal reach and saccade networks was observed. Taken together, this suggests earlier independent effector planning for reaches than saccades, with the saccade network becoming fully engaged around the time of movement execution. The second question we aimed to answer was how saccade and reach differ from each other during the planning and execution phases. To do this, we directly contrasted saccade and reach activation during these time periods and found that for both planning and execution, the effector-preference activation corresponding to reaches was much stronger than the activation corresponding to saccades. During the planning phase, no region showed greater activation for saccades than reaches, and only right FEF showed greater activation for saccades than reaches during execution. The third question we aimed to answer was how effector-specific movement planning and execution are temporally and spatially distributed through the brain. We found that for the within-effector analysis, the time courses for ROIs during the effector-preference planning and execution phases were very highly correlated across parietal and frontal brain regions for both reach and saccade. When comparing activation across effectors, there were much stronger correlations in the right hemispheres (ipsilateral to the arm use to reach) than the contralateral left hemisphere, where AG was the only area strongly correlated across effectors. This suggests left AG may serve a similar role in right-handed reach planning and saccades, the activation seen in other areas is more heavily modulated depending on the effector to be used.

### General activation during visual target memory, reach planning, and reach execution

Many previous fMRI studies have implicated a widespread number of areas in occipital, parietal, and frontal cortex in visually guided reaching (Astafiev et al., 2003, Connolly et al., 2003; Medendorp et al., 2003, 2005; Prado et al., 2005; Fernandez-Ruiz et al., 2007; Beurze et al., 2009; Cavina-Pratesi et al., 2010; Fabbri et al., 2012; Konen et al., 2013; Chen et al., 2014) and saccades (Astafiev et al., 2003; Beurze et al., 2009; Connolly et al, 2007; Curtis et al, 2008; Gallivan et al, 2011, 2015; Bestman et al., 2012; Herwig et al, 2014). However, to our knowledge none of these studies clearly separated the three phases of effector-agnostic target representation, independent effector movement planning, and independent effector movement execution through time. To do this within the spatiotemporal limitations of fMRI, we required a paradigm with a series of instructions and delays which likely introduced more cognitive aspects to the task than one would see during on-line control, but with this caveat in mind, we were able to trace both general and direction-specific activation through those three phases. Most of our regions of interest showed different degrees of time-locked activation during target representation, planning, and execution (Figures 5 and 6), depending on whether the region was more visual (e.g., SOG) or motor (e.g., PMd/M1), but here we will restrict our discussion to significant clusters of activation during these three phases (Figures 2 and 3).

Our analysis of the effector-agnostic target representation phase (Figure 2) revealed limited activation in bilateral mIPS and left AG, perhaps related to spatial working memory (Courtney et al., 1996; Srimal and Curtis, 2008) or activity related to preparatory set (Culham et al., 2006, Chen et al., 2014). Cappadocia et al. (2017) found that bilateral PMd and right pIPS are involved in the target memory phase for reach planning, and activation in the parietal cortex is consistent with the uncertainty condition found in Gertz and Fiehler (2015), where parietal activation was also in the left hemisphere.

During this movement planning phase (Figure 3A), we observed widespread activation in the medial parieto-frontal reach network for reaches, including SPOC, mIPS, PMd, and left M1 (Culham et al, 2006; Gallivan and Culham, 2015; Cappadocia et al., 2017). This was consistent to the activation previously observed in Cappadocia et al. (2017), and further suggests that previous studies that combined target representation and movement planning (Medendorp et al., 2003, Fernandez-Ruiz et al., 2007; Culham et al, 2011) were mainly reporting activity related to visuomotor transformations and/or movement planning, as opposed to target memory. The activation we observed for saccades builds on the importance of separating out the target representation and planning phases, as we were able to observe bilateral pIPS activation in the saccade motor network. Previous research has shown the parietal saccade-related areas to be involved in both eye movements and attention (Corbetta et al., 1998; Astafiev et al., 2003), eye movements and visual working memory (Curtis et al, 2004; Curtis and Connelly, 2008; Srimal and Curtis, 2008) and all three processes (Jerde et al., 2012), and it is very possible that the activation we observed incorporates both early motor planning for saccades, as well as a memory component.

We also observed activation of occipital cortex, including LG, IOG and SOG, during the movement planning phase for both reaches and saccades. A phenomenon known as ‘occipital reactivation’ (Singhal et al., 2013), which involves re-entrant feedback from motor systems and has been previously observed for reaches (Singhal et al., 2013; Cappadocia et al., 2017), could explain this activation. For reaches, lateral cortex activation was greater in the left hemisphere contralateral to the hand, consistent with previous studies (Connolly et al., 2003; Fernandez-Ruiz, 2007; Bernier et al., 2012; Gertz and Fiehler, 2015; Cappadocia et al., 2017).

During motor execution, all medial reach-related regions of activation noted in the planning phase became even more extensive (relative to controls) (Figure 3B), also extending into prefrontal (e.g., SFG) and inferior parietal (e.g., SMG) areas that might be associated with cognitive aspects of the task, such as guidance of the movement based on spatial memory (Gallivan et al., 2015, Cappadocia et al., 2017). For saccade trials, the more lateral parieto-frontal saccade network became much more engaged at the time of execution, including bilateral activation of the FEF, mIPS, and AG. This activation is consistent with saccade-related activity observed in other studies (Astafiev et al., 2003; Beurze et al., 2009; Connolly et al, 2007; Curtis et al, 2008; Gallivan et al, 2011, 2015; Bestman et al., 2012; Herwig et al, 2014), and suggests that the saccade system becomes more heavily engaged much closer to the time of execution than the reach network. Given the nature and frequency of saccades, which are preformed 3-5 times per second (Rayner et al., 1998), this finding is not surprising.

The conjunction analysis identifying areas involved in both reach and saccade execution (figure 4) revealed left mIPS and right AG, PCu, SPOC, LOTC, and FEF/PMv as active for both reaches and saccades (compared to the colour control task). The inclusion of both parietal and frontal motor areas in planning for both effectors is consistent with previous studies, and may explain some of the difficulties in observing effector-preferential activation (Connolly et al, 2007; Curtis et al, 2008; Beurze et al., 2009; Gallivan et al, 2011). The increased overlap in the right hemisphere can likely be explained by the overall increase in activation observed for the saccade>colour analysis, and may not necessarily be reflective of true increased overlap in the right hemisphere. This could be due to the motor requirements of the control task (which required the subject to press a button once or twice with their right hand indicating if the target was red or green) increasing the baseline control activation in the left hemisphere and thus limiting the number of saccade regions that could be identified by the contrast.

The later activation of the saccade network compared to the reach network is further supported when examining the time courses for saccade and reach execution ROIs in figures 5 and 6. In both sets of time courses, most of the ROIs show significantly higher activation when compared to the control condition, with left SOG, left mIPS, and left SPOC showing significantly higher activation during execution for both effectors. As mIPS and SPOC have been implicated in both reach and saccade planning (Culham et al., 2006; Gallivan et al., 2011, Cappadocia et al., 2017) it is not surprising that these areas show significant execution-related activation for both effectors. Although we did not run statistics on this, one can also observe that for reach, bilateral mIPS, SPOC, and PMd all show activation above the control condition during the planning phase, while only left SPOC shows this for saccades. This observationally supports our finding that the saccade network becomes engaged later than the reach network during the execution phase.

### Effector Preference during Movement Planning and Execution

A second goal of our study was to look at cortical effector preference during movement planning and execution. To do this, we contrasted reach and saccade task activation during both movement planning and execution. During the planning phase, we found only effector-preference activation for reach. This is consistent with previous studies that have seen fMRI activation related to reaches produce broader and more spread out BOLD activation than that seen for saccades (DeSouza et al. 2000; Astafiev et al., 2003; Beurze et al., 2007; Gallivan et al., 2011). A similar trend was observed during the motor execution phase, with right FEF appearing as the sole saccade effector-selective ROI. This explanation is further supported by the time course analysis shown in figure 8, where preferential activation can be observed for reaching during the movement planning phase, and even more so during execution. In fact, bilateral PMd, bilateral mIPS, bilateral SMG, and left M1 all showed significantly higher % signal change during execution, even though for all areas but left M1, there was clear activation observed for saccades as well. The complete lack of saccade-related areas in the left hemisphere during the planning phase (figure 7A) may be due to contralateral direction tuning and handedness for reaches that has been observed in other studies: fMRI (Medendorp et al. 2003; 2005; Filimon, 2010; Vesia and Crawford, 2012; Gertz and Fiehler, 2015; Cappadocia et al., 2017), MEG (Van Der Werf et al., 2010), TMS (Vesia et al. 2010), patients (Khan et al., 2007) and primate neurophysiology (Gail and Andersen, 2006; Gail et al., 2009; Westendorff et al., 2010). This asymmetry could also relate to the statics of fMRI, i.e., the way that several neural signals might need to combine to produce significant effects at the level of the BOLD signal. It is also possible that we observed reach-preferential planning due to the increased complexity required for reaches (more degrees of freedom, complex movement dynamics, and the need for hand position information; see Blohm et al., 2007, 2008), thus requiring more planning before the initiation of a movement. It is also possible that there was some implicit saccade planning in the reach task, as eye and arm movements are normally coupled (Flanagan and Johansson, 2003).

### Shared inputs within and between effector-preference networks

To understand saccade and reach planning at a network level, we cross-correlated saccade activation for ROIs observed during reach and saccade motor execution within and across effectors. We found that within each effector, the time courses for ROIs during the planning and execution phases were very highly correlated across parietal and frontal brain regions. For reaches (Figure 9C/D), there were significant correlations between almost all pairs of areas in the right hemisphere (SPOC, mIPS, PCu, AG, SMG, PMv, and PMd), with the one exception being the pairing of SPOC and PMd. In the left hemisphere, significant correlations were found between mIPS, aIPS, AG, SMG, PMv, PMd, and M1 (with the two exceptions being SMG-mIPS and SMG-aIPS). Left SPOC, however, did not significantly correlate with any of the other areas in the left hemisphere. Recent studies have implicated SPOC as a visually-guided reaching area (Culham et al., 2006; Filimon et al., 2009; Vesia et al., 2010; Gallivan and Culham, 2015, Cappadocia et al., 2017), and our previous study showed that for reach, SPOC was identified as a ‘hub’ areas, showing both visual and movement selective activation (Cappadocia et al., 2017). Its important to note that our previous study was examining visual and motor selectivity, and on the visual selectivity index (which is similar to trials in this study), SPOC was only significantly correlated to 5/10 reach-related areas in the left hemisphere.

For saccades (Figure 9A/B), in both the left and right hemisphere mIPS and AG appear as ‘hubs’ that are highly correlated with the activation seen in several other areas. mIPS specifically was significantly correlated with all areas in the right hemisphere and 4/5 other areas in the left hemisphere (the exception being SPOC). Given the strong research linking mIPS to saccade planning (Astafiev et al., 2003; Beurze et al., 2009; Connolly et al, 2007; Curtis et al, 2008; Gallivan et al, 2011, 2015; Bestman et al., 2012; Herwig et al, 2014), it is not surprizing that mIPS was observed to be a highly correlated ‘hub’. AG was previously found to be visually selective for reach planning (Cappadocia et al., 2017) and is involved in left/right spatial discrimination (Hirnstein et al., 2011). It has also been implicated as a saccade planning region in previous TMS tasks (Vesia et al., 2010). Previous TMS studies targeting P3/P4 [which is around the angular gyrus (Ryan et al., 2006)] have found various saccade effects (Elkington et al., 1992; Oyachi and Ohtsuka, 1995; Muri et al., 1996; Kapoula et al., 2005), and it has been suggested that AG is a human homologue of monkey LIP (Koyama et al., 2004), which support its inclusion as a ‘hub’.

Comparing across effectors (Figure 10), there were much stronger correlations in the right hemisphere than in the left. As mentioned earlier in this discussion, this could be due to the high degree of contralateral activation seen for reaches when compared with ipsilateral activation. This generally agrees with previous investigations of occipital, parietal, and prefrontal activity based on fMRI (Medendorp et al. 2003; 2005; Filimon, 2010; Vesia and Crawford, 2012; Gertz and Fiehler, 2015; Cappadocia et al., 2017), MEG (Van Der Werf et al., 2010), TMS (Vesia et al. 2010), patients (Khan et al., 2007) and primate neurophysiology (Gail and Andersen, 2006; Gail et al., 2009; Westendorff et al., 2010). Bilaterally, AG was found to be a ‘hub’ area, showing high correlations across effectors. This is consistent with previous TMS studies to the nearby P3/P4 sights, which have shown that TMS to AG disrupts the integration of saccade information for reach planning (Van Donkelaar et al., 2000). AG has also shown to code motor coordinates, perhaps in somatosensory coordinates (Fernandez-Ruiz et al., 2007; Vesia et al., 2006, 2010; Vesia and Crawford, 2012), signaling it may integrate visual information to produce a motor output common to both effectors. Within the left hemisphere, SPOC, FEF, and SFG from the saccade contrast all showed minimal significant correlations to reach areas (SPOC was only correlated to the reach-SPOC and SFG was only correlated to aIPS). This suggests that these areas are more effector-specific, with recent primate neurophysiology evidence implicating FEF as transitioning from a more general visual code to a saccade movement code (Sajad et al., 2016). While this has not been assessed for effector specificity, the role of the FEF in saccade target selection and production is consistent with its output being more effector-specific (Andersen et al. 1985; Bruce et al., 1985; Schall and Hanes, 1993; Schall et al., 1995; Tehovnik et al., 2000).

### Implications for independent effector control and eye-hand coordination

Consistent with previous studies, there was considerable overlap observed for both reaches and saccades around the time of movement execution (Connolly et al, 2007; Curtis et al, 2008; Beurze et al., 2009; Gallivan et al, 2011). When the timeseries were analyzed between reaches and saccades, an interesting pattern emerged between the hemispheres. As observed in figure 10, almost all right hemisphere brain areas (37/42) show statistically significant correlations, compared to the left hemisphere where only 15/48 regions (31%) are significantly correlated. As all subjects were right handed and reached with their dominant right arm, this could be related to increased reach activation in the left hemisphere (Medendorp et al. 2003; 2005; Filimon, 2010; Vesia and Crawford, 2012; Gertz and Fiehler, 2015; Cappadocia et al., 2017). However, the strong overlap in brain areas revealed in the effector-dependent contrasts combined with the strong correlations observed between reach and saccade areas at execution suggests that for independent or coordinated movements using the eye and hand, a similar network is activated.

## Conclusion

This study shows that when separating visually guided reaches and saccades into distinct target memory, movement planning, and movement execution phases, the evolution of these processes over time reveal increased activation. During effector-agnostic target memory, only modest activation in left AG and bilateral mIPS was observed. For reaches, the occipital-parietal-frontal reach network was engaged medially during both reach *planning and execution*. For saccades, more lateral (mIPS, AG, and FEF) activity was observed only during saccade *execution*. These motor activations were bilateral, with a left (contralateral) preference for reach. With the exception of right FEF, effector-preference contrasts revealed significantly more parietofrontal activation for reaches than saccades during both planning and execution. Cross-correlation of reach, saccade, and reach-saccade activation through time revealed spatiotemporally correlated activation both within and across effectors in each hemisphere, but with a tendency toward higher correlations in the right hemisphere. These results demonstrate cortical networks for eye, hand, and eye-hand control that are widely distributed, but with effector-specific rules for timing, medial-lateral localization, and hemispheric lateralization.

## References

Astafiev S V, Shulman GL, Stanley CM, Snyder AZ, Van Essen DC, Corbetta M. 2003. Functional organization of human intraparietal and frontal cortex for attending, looking, and pointing. J Neurosci. 23:4689–4699.

Barany DA, Shapiro AD, Lee TG. (2015) Multivariate fMRI Approaches to Flexible Sensorimotor Maps in Parietal Cortex. J. Neurosci. 35(34):11763–11765

Batista, A. P., Buneo, C. A., Snyder, L. H., & Andersen, R. A. (1999). Reach plans in eye-centered coordinates. Science, 285(5425), 257–260. doi: 10.1126/science.285.5425.257

Battaglia-Mayer, A., Ferraina, S., Mitsuda, T., Marconi, B., Genovesio, A., Onorati, P.,.. Caminiti, R. (2000). Early coding of reaching in the parietooccipital cortex. Journal of Neurophysiology, 83(4), 2374–2391.

Bernier P-M, Cieslak M, Grafton ST. 2012. Effector selection precedes reach planning in the dorsal parietofrontal cortex. J Neurophysiol. 108:57–68.

Bernier P-M, Grafton ST. 2010. Human posterior parietal cortex flexibly determines reference frames for reaching based on sensory context. Neuron. 68:776–788.

Bernier P-M, Whittingstall K,Grafton ST. 2017. Differential Recruitment of Parietal Cortex during Spatial and Non-spatial Reach Planning. Front Hum Neurosci. 2017 11:249.

Beurze SM, de Lange FP, Toni I, Medendorp WP. 2007. Integration of target and effector information in the human brain during reach planning. J Neurophysiol. 97:188–199.

Beurze SM, de Lange FP, Toni I, Medendorp WP. 2009. Spatial and effector processing in the human parietofrontal network for reaches and saccades. J Neurophysiol. 101:3053–3062.

Beurze SM, Toni I, Pisella L, Medendorp WP. 2010. Reference frames for reach planning in human parietofrontal cortex. J Neurophysiol. 104:1736–1745.

Binkofski, F., Dohle, C., Posse, S., Stephan, K. M., Hefter, H., Seitz, R. J., & Freund, H. J. (1998). Human anterior intraparietal area subserves prehension - A combined lesion and functional MRI activation study. Neurology, 50(5), 1253–1259.

Blangero A, Gaveau V, Luauté J, Rode G, Salemme R, Guinard M, Boisson D, Rossetti Y, Pisella L. 2008. A hand and a field effect in on-line motor control in unilateral optic ataxia. Cortex. 44(5):560–568.

Blohm, G., & Crawford, J. D. (2007). Computations for geometrically accurate visually guided reaching in 3-D space. Journal of Vision, 7(5), 4. doi: 10.1167/7.5.4

Blohm, G., & Crawford, J. D. (2009). Fields of gain in the brain. Neuron, 64(5), 598–600. doi: 10.1016/j.neuron.2009.11.022

Buneo, C. A., Jarvis, M. R., Batista, A. P., & Andersen, R. A. (2002). Direct visuomotor transformations for reaching. Nature, 416(6881), 632–636. doi: 10.1038/416632a

Cappadocia DC, Monaco S, Chen Y, Blohm G, Crawford JD. 2017. Temporal Evolution of Target Representation, Movement Direction Planning, and Reach Execution in Occipital–Parietal–Frontal Cortex: An fMRI Study. Cerebral Cortex. 27(11):5242–5260.

Cattaneo, Z., Silvanto, J., Pascual-Leone, A., & Battelli, L. (2009). The role of the angular gyrus in the modulation of visuospatial attention by the mental number line. Neuroimage, 44(2), 563–568. doi: 10.1016/j.neuroimage.2008.09.003

Cavina-Pratesi, C., Valyear, K. F., Culham, J. C., Kohler, S., Obhi, S. S., Marzi, C. A., & Goodale, M. A. (2006). Dissociating arbitrary stimulus-response mapping from movement planning during preparatory period: Evidence from event-related functional magnetic resonance imaging. Journal of Neuroscience, 26(10), 2704– 2713.

Cavina-Pratesi C, Monaco S, Fattori P, Galletti C, McAdam TD, Quinlan DJ, Goodale MA, Culham JC. 2010. Functional magnetic resonance imaging reveals the neural substrates of arm transport and grip formation in reach-to-grasp actions in humans. J Neurosci. 30:10306–10323.

Chen Y, Monaco S, Byrne P, Yan X, Henriques DYP, Crawford JD. 2014. Allocentric versus egocentric representation of remembered reach targets in human cortex. J Neurosci. 34:12515–12526.

Chapman, C. S., Gallivan, J. P., Culham, J. C., & Goodale, M. A. (2011). Mental blocks: FMRI reveals top-down modulation of early visual cortex when obstacles interfere with grasp planning. Neuropsychologia, 49(7), 1703–1717. doi: 10.1016/j.neuropsychologia.2011.02.048

Cisek P, Kalaska JF. 2002. Simultaneous encoding of multiple potential reach directions in dorsal premotor cortex. J Neurophysiol. 87:1149–1154.

Cisek P, Kalaska JF. 2005. Neural correlates of reaching decisions in dorsal premotor cortex: specification of multiple direction choices and final selection of action. Neuron. 45:801–814.

Connolly JD, Andersen RA, Goodale MA. 2003. FMRI evidence for a “parietal reach region” in the human brain. Exp brain Res. 153:140–145.

Connolly JD, Goodale MA, Cant JS, Munoz DP. 2007. Effector-specific fields for motor preparation in the human frontal cortex. Neuroimage. 34:1209–1219.

Connolly JD, Goodale MA, DeSouza JF, Menon RS, Vilis T. 2000. A comparison of frontoparietal fMRI activation during anti-saccades and anti-pointing. J Neurophysiol. 84:1645–1655.

Corbetta, M., Akbudak, E., Conturo, T. E., Snyder, A. Z., Ollinger, J. M., Drury, H. A., Shulman, G. L. (1998). A common network of functional areas for attention and eye movements. Neuron, 21(4), 761–773.

Crawford JD, Henriques DY, Medendorp WP. 2011. Three-dimensional transformations for goal-directed action. Annu Rev Neurosci. 34:309–31.

Culham, J. C., Danckert, S. L., DeSouza, J. F. X., Gati, J. S., Menon, R. S., & Goodale, M. A. (2003). Visually guided grasping produces fMRI activation in dorsal but not ventral stream brain areas. Experimental Brain Research, 153(2), 180–189. doi: 10.1007/s00221-003-1591-5

Culham, J. C., & Kanwisher, N. G. (2001). Neuroimaging of cognitive functions in human parietal cortex. Current Opinion in Neurobiology, 11(2), 157–163. doi: 10.1016/S0959-4388(00)00191-4

Culham, J. C., & Valyear, K. F. (2006). Human parietal cortex in action. Current Opinion in Neurobiology, 16(2), 205–212. doi: 10.1016/j.conb.2006.03.005

Culham, J. C., Cavina-Pratesi, C., & Singhal, A. (2006). The role of parietal cortex in visuomotor control: What have we learned from neuroimaging? Neuropsychologia, 44(13), 2668–2684. doi: 10.1016/j.neuropsychologia.2005.11.003

Curtis CE, Connolly JD. 2008. Saccade preparation signals in the human frontal and parietal cortices. J Neurophysiol. 99(1):133–145.

Curtis CE, Rao VY, D’Esposito M. 2004. Maintenance of spatial and motor codes during oculomotor delayed response tasks. J Neurosci. 24(16):3944–3952.

Curtis CE. 2006. Prefrontal and parietal contributions to spatial working memory. Neuroscience 139(1):173–80.

Desmurget, M., Epstein, C. M., Turner, R. S., Prablanc, C., Alexander, G. E., & Grafton, S. T. (1999). Role of the posterior parietal cortex in updating reaching movements to a visual target. Nature Neuroscience, 2(6), 563–567.

Desmurget M, Pélisson D, Rossetti Y, Prablanc C. 1998. From eye to hand: planning goal-directed movements. Neurosci Biobehav Rev. 22(6):761–88.

DeSouza JF, Dukelow SP, Gati JS, Menon RS, Andersen RA, Vilis T. 2000. Eye position signal modulates a human parietal pointing region during memory-guided movements. J Neurosci. 20:5835–5840.

Dessing, J. C., Vesia, M., & Crawford, J. D. (2013). The role of areas MT+/V5 and SPOC in spatial and temporal control of manual interception: An rTMS study. Frontiers in Behavioral Neuroscience, 7, 15.

Engel SA, Glover GH, Wandell BA. 1997. Retinotopic organization in human visual cortex and the spatial precision of functional MRI. Cereb Cortex. 7:181–192.

Farrer C, Frey SH, Van Horn JD, Tunik E, Turk D, Inati S, Grafton ST. 2008. The angular gyrus computes action awareness representations. Cereb Cortex. 18(2):254–61.

Fattori, P., Gamberini, M., Kutz, D. F., & Galletti, C. (2001). ’Arm-reaching’ neurons in the parietal area V6A of the macaque monkey. European Journal of Neuroscience, 13(12), 2309–2313.

Fattori, P., Breveglieri, R., Marzocchi, N., Filippini, D., Bosco, A., & Galletti, C. (2009). Hand orientation during reach-to-grasp movements modulates neuronal activity in the medial posterior parietal area V6A. Journal of Neuroscience, 29(6), 1928–1936.

Fernandez-Ruiz J, Goltz HC, DeSouza JFX, Vilis T, Crawford JD. 2007. Human parietal “reach region” primarily encodes intrinsic visual direction, not extrinsic movement direction, in a visual motor dissociation task. Cereb Cortex. 17:2283–2292.

Ferraina, S., Battaglia-Mayer, A., Genovesio, A., Marconi, B., Onorati, P., & Caminiti, R. (2001). Early coding of visuomanual coordination during reaching in parietal area PEc. Journal of Neurophysiology, 85(1), 462–467.

Filimon, F. (2010). Human cortical control of hand movements: Parietofrontal networks for reaching, grasping, and pointing. Neuroscientist, 16(4), 388–407.

Filimon, F., Nelson, J. D., Huang, R., & Sereno, M. I. (2009). Multiple parietal reach regions in humans: Cortical representations for visual and proprioceptive feedback during on-line reaching. Journal of Neuroscience, 29(9), 2961–2971.

Flanagan JR, Johansson RS. 2003. Action plans used in action observation. Nature. 424:769–771.

Gallivan JP, Cavina-Pratesi C, Culham JC. 2009. Is that within reach? fMRI reveals that the human superior parieto-occipital cortex encodes objects reachable by the hand. J Neurosci. 29:4381–4391.

Gallivan JP, Culham JC. 2015. Neural coding within human brain areas involved in actions. Curr Opin Neurobiol. 33:141–149.

Gallivan JP, McLean A, Culham JC. 2011. Neuroimaging reveals enhanced activation in a reach-selective brain area for objects located within participants’ typical hand workspaces. Neuropsychologia. 49:3710–3721.

Gertz H, Fiehler K. 2015. Human posterior parietal cortex encodes the movement goal in a pro-/anti-reach task. J Neurophysiol. 114:170–183.

Goodale MA. 2011 Transforming Vision into Action. Vision Res. 51(13):1567-87.

Goodale, M. A., & Milner, A. D. (1992). Separate visual pathways for perception and action. Trends in Neurosciences, 15(1), 20–25.

Goodale, M. A., Milner, A. D., Jakobson, L. S., & Carey, D. P. (1991). A neurological dissociation between perceiving objects and grasping them. Nature, 349(6305), 154–156.

Gorbet, D. J., Staines, W. R., & Sergio, L. E. (2004). Brain mechanisms for preparing increasingly complex sensory to motor transformations. Neuroimage, 23(3), 1100-1111. doi: 10.1016/j.neuroimage.2004.07.043

Gottlieb, J., & Goldberg, M. E. (1999). Activity of neurons in the lateral intraparietal area of the monkey during an antisaccade task. Nature Neuroscience, 2(10), 906–912. doi: 10.1038/13209

Gottlieb, J. P., Kusunoki, M., & Goldberg, M. E. (1998). The representation of visual salience in monkey parietal cortex. Nature, 391(6666), 481–484.

Haar S, Donchin O, Dinstein I (2015) Dissociating visual and motor directional selectivity using visuomotor adaptation. J Neurosci 35:6813–6821.

Hawkins KM, Sayegh P, Yan X, Crawford JD, Sergio LE. 2013. Neural activity in superior parietal cortex during rule-based visual-motor transformations. J Cogn Neurosci. 25:436–454.

Heed, T., Beurze, S. M., Toni, I., Roder, B., & Medendorp, W. P. (2011). Functional rather than effector-specific organization of human posterior parietal cortex. Journal of Neuroscience, 31(8), 3066–3076.

Henriques DY, Klier EM, Smith MA, Lowy D, Crawford JD. 1998. Gaze-centered remapping of remembered visual space in an open-loop pointing task. J Neurosci. 18:1583–1594.

Jerde TA, Merriam EP, Riggall AC, Hedges JH, Curtis CE. 2012. Prioritized maps of space in human frontoparietal cortex. 32(48):17382–90.

Johnson, P. B., Ferraina, S., Bianchi, L., & Caminiti, R. (1996). Cortical networks for visual reaching: Physiological and anatomical organization of frontal and parietal lobe arm regions. Cerebral Cortex, 6(2), 102–119.

Kalaska, J. F., Scott, S. H., Cisek, P., & Sergio, L. E. (1997). Cortical control of reaching movements. Current Opinion in Neurobiology, 7(6), 849–859.

Kapoula, Z., Yang, Q., Coubard, O., Daunys, G., & Orssaud, C. (2005). Role of the posterior parietal cortex in the initiation of saccades and vergence: Right/left functional asymmetry. Clinical and Basic Oculomotor Research: In Honor of David S.Zee, 1039, 184–197.

Konen CS, Mruczek REB, Montoya JL, Kastner S. 2013. Functional organization of human posterior parietal cortex: grasping- and reaching-related activations relative to topographically organized cortex. J Neurophysiol. 109:2897–2908.

Koyama, M., Hasegawa, I., Osada, T., Adachi, Y., Nakahara, K., & Miyashita, Y. (2004). Functional magnetic resonance imaging of macaque monkeys performing visually guided saccade tasks: Comparison of cortical eye fields with humans. Neuron, 41(5), 795–807.

Malik P, Dessing JC, Crawford JD. 2015. Role of early visual cortex in trans-saccadic memory of object features. J Vis. 15(11):7.

Medendorp WP, Goltz HC, Crawford JD, Vilis T. 2005. Integration of target and effector information in human posterior parietal cortex for the planning of action. J Neurophysiol. 93:954–962.

Medendorp WP, Goltz HC, Vilis T. 2006. Directional selectivity of BOLD activity in human posterior parietal cortex for memory-guided double-step saccades. J Neurophysiol. 95:1645–1655.

Medendorp WP, Goltz HC, Vilis T, Crawford JD. 2003. Gaze-centered updating of visual space in human parietal cortex. J Neurosci. 23:6209–6214.

Medendorp, W. P., Buchholz, V. N., Van Der Werf, J., & Leone, F. T. M. (2011). Parietofrontal circuits in goal-oriented behaviour. European Journal of Neuroscience, 33(11), 2017–2027.

Merriam, E. P., Genovese, C. R., & Colby, C. L. (2003). Spatial updating in human parietal cortex. Neuron, 39(2), 361–373.

Miller, L., Sun, F., Curtis, C., & D’Esposito, M. (2005). Functional interactions between oculomotor regions during prosaccades and antisaccades. Human Brain Mapping, 26(2), 119–127.

Milner, A. D., Dijkerman, H. C., Pisella, L., McIntosh, R. D., Tilikete, C., Vighetto, A., & Rossetti, Y. (2001). Grasping the past: Delay can improve visuomeotor performance. Current Biology, 11(23), 1896–1901.

Monaco S, Cavina-Pratesi C, Sedda A, Fattori P, Galletti C, Culham JC. 2011. Functional magnetic resonance adaptation reveals the involvement of the dorsomedial stream in hand orientation for grasping. J Neurophysiol. 106:2248–2263.

Olson CR. 2003. Brain representation of object-centered space in monkeys and humans. Annu Rev Neurosci. 26:331–354.

Petit, L., & Haxby, J. V. (1999). Functional anatomy of pursuit eye movements in humans as revealed by fMRI. Journal of Neurophysiology, 82(1), 463–471.

Picard N, Strick PL. 2001. Imaging the premotor areas. Curr Opin Neurobiol. 11:663–672.

Pisella, L., Sergio, L., Blangero, A., Torchin, H., Vighetto, A., & Rossetti, Y. (2009). Optic ataxia and the function of the dorsal stream: Contributions to perception and action. Neuropsychologia, 47(14), 3033–3044.

Prablanc, C., Echallier, J. F., Komilis, E., & Jeannerod, M. (1979). Optimal response of eye and hand motor systems in pointing at a visual target. 1. spatio-temporal characteristics of eye and hand movements and their relationships when varying the amount of visual information. Biological Cybernetics, 35(2), 113–124.

Prado J, Clavagnier S, Otzenberger H, Scheiber C, Kennedy H, Perenin M-T. 2005. Two cortical systems for reaching in central and peripheral vision. Neuron. 48:849–858.

Prime, S. L., M. Vesia, et al. (2008). Transcranial magnetic stimulation over posterior parietal cortex disrupts transsaccadic memory of multiple objects. Journal of Neuroscience 28(27): 6938–6949.

Prime, S. L., M. Vesia, et al. (2010). TMS Over Human Frontal Eye Fields Disrupts Trans-saccadic Memory of Multiple Objects. Cerebral Cortex 20(4): 759–772.

Quinlan, D. J., & Culham, J. C. (2007). fMRI reveals a preference for near viewing in the human parieto-occipital cortex. Neuroimage, 36(1), 167–187.

Reed CL, Grubb JD, Steele C. 2006. Hands up: attentional prioritization of space near the hand. J Exp Psychol Hum Percept Perform. 32:166–177.

Rice, N. J., Tunik, E., Cross, E. S., & Grafton, S. T. (2007). On-line grasp control is mediated by the contralateral hemisphere. Brain Research, 1175, 76-84.

Rice, N. J., Tunik, E., & Grafton, S. T. (2006). The anterior intraparietal sulcus mediates grasp execution, independent of requirement to update: New insights from transcranial magnetic stimulation. Journal of Neuroscience, 26(31), 8176–8182.

Rizzolatti, G., Fadiga, L., Gallese, V., & Fogassi, L. (1996). Premotor cortex and the recognition of motor actions. Cognitive Brain Research, 3(2), 131–141.

Rizzolatti, G., & Luppino, G. (2001). The cortical motor system. Neuron, 31(6), 889-901.

Rossetti Y, Pisella L, Vighetto A. 2003. Optic ataxia revisited: visually guided action versus immediate visuomotor control. Exp Brain Res. 153(2):171–179.

Sadeh M, Sajad A, Wang H, Yan X, Crawford JD. 2015. Spatial transformations between superior colliculus visual and motor response fields during head-unrestrained gaze shifts. Eur J Neurosci. 42(11):2934–51.

Sajad A, Sadeh M, Keith GP, Yan X, Wang H, Crawford JD. 2015. Visual-Motor Transformations Within Frontal Eye Fields During Head-Unrestrained Gaze Shifts in the Monkey. Cereb Cortex. 25(10):3932–52.

Sajad A, Sadeh M, Yan X, Wang H, Crawford JD. 2016. Transition from Target to Gaze Coding in Primate Frontal Eye Field during Memory Delay and Memory–Motor Transformation. e0040-16.2016 1–20.

Schluppeck, D., Curtis, C., Glimcher, P., & Heeger, D. (2006). Sustained activity in topographic areas of human posterior parietal cortex during memory-guided saccades. Journal of Neuroscience, 26(19), 5098–5108.

Schluppeck, D., Glimcher, P., & Heeger, D. J. (2005). Topographic organization for delayed saccades in human posterior parietal cortex. Journal of Neurophysiology, 94(2), 1372–1384.

Singh KD, Smith AT, Greenlee MW. 2000. Spatiotemporal frequency and direction sensitivities of human visual areas measured using fMRI. Neuroimage. 12:550–564.

Singhal A, Monaco S, Kaufman LD, Culham JC. 2013. Human fMRI reveals that delayed action re-recruits visual perception. PLoS One. 8(9):e73629.

Smith, E., & Jonides, J. (1998). Neuroimaging analyses of human working memory. Proceedings of the National Academy of Sciences of the United States of America, 95(20), 12061–12068. doi: 10.1073/pnas.95.20.12061

Srimal R, Curtis CE. 2008. Persistent neural activity during the maintenance of spatial position in working memory. Neuroimage. 39(1):455–468.

Tosoni A, Galati G, Romani GL, Corbetta M. 2008. Sensory-motor mechanisms in human parietal cortex underlie arbitrary visual decisions. Nat Neurosci. 11:1446–1453.

Valyear, K. F., Gallivan, J. P., McLean, D. A., & Culham, J. C. (2012). fMRI repetition suppression for familiar but not arbitrary actions with tools. Journal of Neuroscience, 32(12), 4247–4259.

Van Der Werf J, Jensen O, Fries P, Medendorp WP. 2008. Gamma-band activity in human posterior parietal cortex encodes the motor goal during delayed prosaccades and antisaccades. J Neurosci. 28(34):8397–8405.

Van Der Werf J, Jensen O, Fries P, Medendorp WP. 2010. Neuronal synchronization in human posterior parietal cortex during reach planning. J Neurosci. 30(4):1402– 1412.

Vesia M, Crawford JD. 2012. Specialization of reach function in human posterior parietal cortex. Exp brain Res. 221:1–18.

Vesia M, Monteon JA, Sergio LE, Crawford JD. 2006. Hemispheric asymmetry in memory-guided pointing during single-pulse transcranial magnetic stimulation of human parietal cortex. J Neurophysiol. 96(6):3016–3027.

Vesia M, Prime SL, Yan X, Sergio LE, Crawford JD. 2010. Specificity of human parietal saccade and reach regions during transcranial magnetic stimulation. J Neurosci. 30:13053–13065.

Vesia, M., Yan, X., Henriques, D. Y., Sergio, L. E., & Crawford, J. D. (2008). Transcranial magnetic stimulation over human dorsal-lateral posterior parietal cortex disrupts integration of hand position signals into the reach plan. Journal of Neurophysiology, 100(4), 2005–2014.

Westendorff S, Klaes C, Gail A. 2010. The cortical timeline for deciding on reach motor goals. J Neurosci. 30:5426–5436.

Wise, S. P., Boussaoud, D., Johnson, P. B., & Caminiti, R. (1997). Premotor and parietal cortex: Corticocortical connectivity and combinatorial computations. Annual Review of Neuroscience, 20, 25–42.

Zhang M, Barash S. 2000. Neuronal switching of sensorimotor transformations for antisaccades. Nature. 408(6815):971–975.

